# Altruistic feeding and cell-cell signaling during bacterial differentiation actively enhance phenotypic heterogeneity

**DOI:** 10.1101/2024.03.27.587046

**Authors:** Taylor B. Updegrove, Thomas Delerue, Vivek Anantharaman, Hyomoon Cho, Carissa Chan, Thomas Nipper, Hyoyoung Choo-Wosoba, Lisa M. Jenkins, Lixia Zhang, Yijun Su, Hari Shroff, Jiji Chen, Carole A. Bewley, L. Aravind, Kumaran S. Ramamurthi

## Abstract

Starvation triggers bacterial spore formation, a committed differentiation program that transforms a vegetative cell into a dormant spore. Cells in a population enter sporulation non-uniformly to secure against the possibility that favorable growth conditions, which puts sporulation-committed cells at a disadvantage, may resume. This heterogeneous behavior is initiated by a passive mechanism: stochastic activation of a master transcriptional regulator. Here, we identify a cell-cell communication pathway that actively promotes phenotypic heterogeneity, wherein *Bacillus subtilis* cells that start sporulating early utilize a calcineurin-like phosphoesterase to release glycerol, which simultaneously acts as a signaling molecule and a nutrient to delay non-sporulating cells from entering sporulation. This produced a more diverse population that was better poised to exploit a sudden influx of nutrients compared to those generating heterogeneity via stochastic gene expression alone. Although conflict systems are prevalent among microbes, genetically encoded cooperative behavior in unicellular organisms can evidently also boost inclusive fitness.

## INTRODUCTION

Genetically identical populations of cells generally behave similarly, especially during robust growth conditions in a uniform environment. However, at lower cell densities and when gene expression is reduced, slight variations in local growth conditions or intracellular protein levels can result in divergent cell behaviors (*1*). This situation, termed “phenotypic heterogeneity”, underlies myriad phenomena such as antibiotic tolerance in bacteria, differences in tumor onset and progression rates, and task allocation in a unicellular population (*2–5*). Phenotypic heterogeneity may also be driven by ephemeral chromosomal changes in a subpopulation of cells (“phase variation”) that result in either differential transcription or the expression of different alleles of a gene in different cells (*6*). A common feature of generating heterogeneity is typically an intrinsically stochastic mechanism that underlies the process. For example, when the intracellular copy number of a master regulator is very low, slight cell-to-cell variations in the production of that factor can vary widely between cells and result in varying phenotypes. Similarly, the unequal partitioning of key regulatory factors between daughter cells can also give rise to heterogeneous behavior. Even in phase variation, the generation of frameshifts, for example, by slipped-strand mispairing during DNA replication at error-prone sites is a stochastic event. Despite the stochastic underpinning, the generation of diversity is thought to be advantageous as it provides a method for a population to poise itself against rapid environmental changes which may otherwise put a homogeneous population at a disadvantage (*7*).

Bacterial endospore formation is a distinctive example of heterogeneity generation during a committed developmental program. When *Bacillus subtilis* encounters starvation, it initiates the sporulation program to transform the normally rod-shaped cell into an ovoid, dormant spore (*8, 9*). Entry into sporulation is governed by phosphorylation of a transcriptional regulator named Spo0A. Once the pool of intracellular phosphorylated Spo0A reaches a high enough threshold, sporulation initiates with the activation of several sporulation-specific genes (*10–12*). An early hallmark of sporulation is the asymmetric division of the rod-shaped cell into two unequal daughter cells: a larger mother cell and a smaller forespore (Fig. 1A). In the next step, the mother cell engulfs the forespore, whereupon the forespore gradually matures into dormancy. Eventually, the mother cell lyses, which releases the now-mature spore into the environment. After asymmetric division, the sporulation program, which takes approximately 6 h to complete, is irreversible, even if there is an influx of nutrients. Thus, cells that commit to sporulate do so at a high risk. If the nutrient limitation that triggers sporulation is transient, sporulation-committed cells are at a disadvantage compared to non-sporulating cells that can take advantage of new nutrients to grow and divide. However, entry into sporulation is naturally asynchronous, which results in a subpopulation of non-sporulating cells that may be available to take advantage of an influx of nutrients while another subpopulation continues to sporulate. This heterogeneity in the population is initially generated by the stochastic phosphorylation of Spo0A (*13, 14*), and the generation of such a subpopulation is thought to provide a bet-hedging strategy against transient environmental changes (*15*).

**Figure 1.**
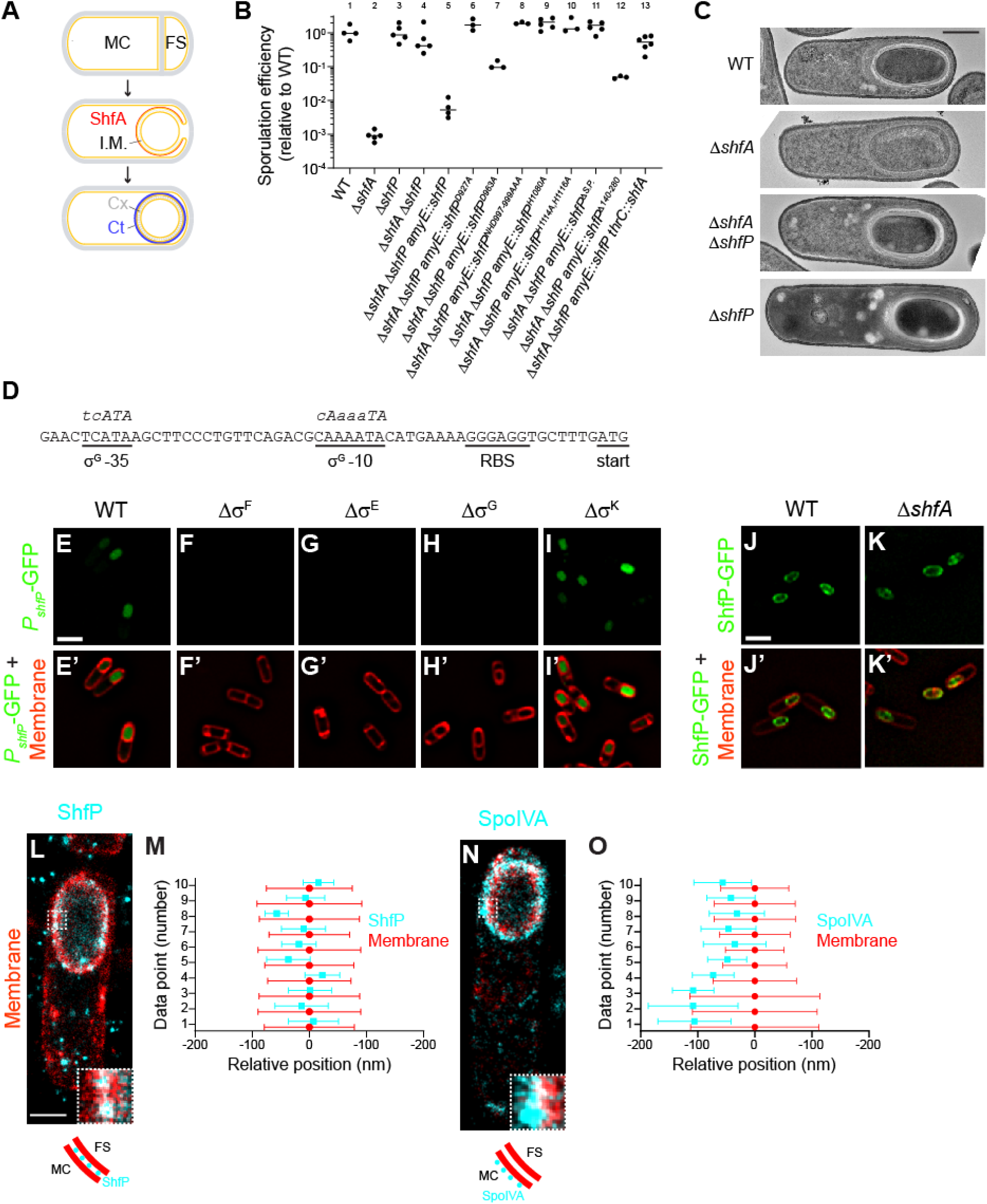
Deletion of *shfP* suppresses the sporulation defect caused by deletion of *shfA*. (A) Schematic of sporulation in *Bacillus subtilis*. Asymmetric division yields two daughter cells (top panel): a larger mother cell (MC) and smaller forespore (FS). Membranes depicted in yellow; cell wall depicted in gray. Middle panel: engulfment of the forespore by the mother cell. ShfA (red) is produced in the mother cell and localizes to the forespore surface. The double membrane envelope produces an intermembrane macromolecular matrix (I.M.) that surrounds the forespore. Bottom panel: the forespore eventually resides as a topologically isolated organelle inside the mother cell and localizes to the forespore surface. The double membrane envelope produces an intermembrane macromolecular matrix (I.M.) that surrounds the forespore. Bottom panel: the forespore eventually resides as a topologically isolated organelle inside the mother cell cytosol; the cortex (Cx, composed of specialized peptidoglycan, depicted in gray dashes), an outer forespore membrane, and the spore coat (a proteinaceous shell, Ct, depicted in blue). (B) Sporulation efficiencies, as measured by resistance to 80°C for 20 min relative to WT, of indicated *B. subtilis* strains. Bars represent mean values; data points represent sporulation efficiencies from an independent culture. Strains: PY79; CW202; TD517; TD507; CC2; CC14; CC15; CC16; CC17; CC18; CC19; CC20; CC232. (C) Electron micrographs of negatively stained thin sections of sporulating wild type, Δ*shfA*, Δ*shfA* Δ*shfP*, or Δ*shfP* strains of *B. subtilis* collected 5 h after induction of sporulation. Scale bar: 500 nm. Strains: PY79; CW202; TD517; TD507. (D) Nucleotide sequence of the region upstream of the *shfP* open reading frame. Putative ribosome binding site (RBS), start codon, and −10 and −35 binding site for σ^G^ are underlined. Consensus nucleotide sequences for −10 and −35 binding sites for σ^G^ are shown in italics above the chromosomal nucleotide sequence, where capital letters depict highly conserved nucleotides. (E-I’) Production (E, I) or lack of production (F-H) of GFP (green) produced under control of *shfP* promoter in (E) wild type (CC182) strain of sporulating *B. subtilis*, or in the absence of (F) σ^F^ (CC188), (G) σ^E^ (CC189), (H) σ^G^ (CC190), or (I) σ^K^ (CC191) 4 h after induction of sporulation. (E’-I’) Overlay of GFP fluorescence (green) in (E-I) and membranes (red). Scale bar: 2 µm. (J-K’) Localization of ShfP-GFP in the presence (J-J’) or absence (K-K’) of ShfA. (J’-K’) Overlay of GFP fluorescence (green) and membranes (red). Scale bar: 2 µm. Strains: CC179; CC175. (L-O) DNA-PAINT super-resolution microscopy of ShfP (L-M) and SpoIVA (N-O) localization to the inter-membrane space and outer forespore membrane surface, respectively. (L, N) ShfP and SpoIVA are colored in cyan, the membrane is colored in red. Portions of each image (white-dashed box) are shown enlarged in the lower right corner of each microscopic image. Below each microscopic image is a carton depiction of the ShfP and SpoIVA (cyan circles) localization relative to the mother cell (MC) and forespore (FS) adjacent membranes (red). Scale bar: 1 µm. Strains: TU46; TU53. (M, O) Quantification of relative localizations of ten 10 different line profiles of ShfP or SpoIVA (cyan) and fluorescence from membrane marker (red). Data point represents the center position for each focus and whiskers are the S.D. representing the width of the distribution of the fluorescence.

Here, we describe a genetically encoded pathway that actively enhances the heterogeneous entry into sporulation in *B. subtilis*. We report that cells that commit early to sporulate are programed to generate and catalytically release glycerol into the environment. We show that the extracellular glycerol functions as a nutrient that delays other cells from entering the sporulation pathway. Additionally, glycerol serves as a cell-cell signaling molecule that represses the buildup of phosphorylated Spo0A and delays entry into sporulation in those cells that have not yet started to sporulate. As a result, wild type (WT) *B. subtilis* generates a more heterogeneous population of cells at different stages of sporulation compared to mutant strains that are not programed to release glycerol. Finally, we show that cells harboring the altruistic feeding and cell-cell communication pathway are better poised to take advantage of an influx of nutrients after the initiation of sporulation compared to cells that generate heterogeneity by stochastic activation of Spo0A alone. We therefore propose that the generation of diversity in a population of differentiating cells can be genetically programed and not simply be a consequence of random events.

## RESULTS

### Deletion of shfP suppresses the sporulation defect caused by shfA deletion

ShfA (<u>S</u>porulation <u>h</u>eterogeneity factor <u>A</u>ntidote; previously YabQ) is produced in the mother cell and localizes to the forespore surface (*16*). It features the sporulating Bacillota-specific three-transmembrane (TM) YabQ domain. This domain has a unique predicted structure with an extracellular N-terminus, conserved intramembrane polar residues, and a kinked third TM helix bracing the first two TM helices. In ShfA proteins, the YabQ domain may be followed by a variable C-terminal extension that in some species contains additional hydrophobic helices (three in the case of *B. subtilis*) that might associate with the membrane. Deletion of *shfA* results in a ∼1000-fold reduction in sporulation efficiency relative to WT (Fig. 1B), presumably due to incomplete elaboration of the cortex peptidoglycan layer (Fig. 1C, panel 2) (*17, 18*). To investigate the function of ShfA, we isolated suppressor mutants that would correct this sporulation defect. Cultures of the Δ*shfA* strain were grown in sporulation media, allowed to accumulate spontaneous mutations, then subjected to repeated cycles of sporulation, followed by exposure to high heat to eliminate cells that were unable to sporulate, followed by germination and re-growth in fresh medium (*19*). Our selection yielded two independent extragenic suppressor mutations that restored sporulation efficiency to near WT levels. Whole genome sequencing revealed that both suppressor mutations mapped to the *yvnB* locus, a gene of previously unknown function which we renamed *shfP* (<u>s</u>porulation <u>h</u>eterogeneity factor <u>P</u>oison) whose transcript is reportedly upregulated late in the sporulation program (*20*). Both mutations (deletion of G3332 and a deletion of residues AAGGA at positions 3529-3533) resulted in frameshifts that introduced a premature stop codon near the 3’ end of the gene. To test if the suppressor mutations resulted in a loss of ShfP function, we examined if deletion of *shfP* would also correct the sporulation defect of the Δ*shfA* mutant. Mutants harboring a deletion of both *shfA* and *shfP* sporulated at near WT levels (Fig. 1B, lane 4) and elaborated a cortex similar to WT (Fig. 1C, panel 3). Complementation of *shfP* at an ectopic chromosomal locus reduced sporulation efficiency to near Δ*shfA* levels (Fig. 1B, lane 5), while additional complementation of this strain with *shfA* again restored sporulation to near WT levels (Fig. 1B, lane 13). In contrast, cells harboring a deletion of *shfP* alone did not exhibit an obvious sporulation defect (Fig. 1B, lane 3) nor did they display an obvious cortex assembly defect (Fig. 1C, panel 4). The data are therefore consistent with a model in which ShfP inhibits sporulation and ShfA counteracts the negative effects of ShfP.

### ShfP is a member of a vast radiation of bacterial cell surface calcineurin-like phosphoesterases

ShfP is a 1289 amino acid secreted protein with an N-terminal signal peptide followed by 9 globular domains (Fig. 2A). The first of these is a Lamin N-terminal domain (LTD), prototyped by the globular DNA-binding and intra-nuclear interaction domain of the animal nuclear envelope lamins (*21*). This is followed by seven immunoglobulin-like (Ig) β-sandwich domains and a calcineurin-like phosphoesterase domain inserted into the seventh Ig domain. Other than nuclear lamins, the LTD was previously reported as fused to different phosphoesterase catalytic domains in several bacterial cell-surface proteins and, based on its animal counterpart, is predicted to bind extracellular DNA or chemically analogous cell-surface polysaccharides (*21*). The Ig domains are similarly predicted to bind cell surface carbohydrates or mediate adhesion via interaction with other proteins. The calcineurin-like domain of ShfP belongs to a vast radiation of such domains in secreted/membrane-anchored proteins that are found across the bacterial superkingdom (Fig. 2A, S1A). These versions are typically distinguished by multiple fusions to carbohydrate-binding and cell-surface adhesion domains which, in addition to LTD and Ig, includes domains such as the concanavalin lectin-like Laminin G, Fibronectin Type III (FN3), S-layer homology (SLH), discoidin-like, cadherin, cell-wall-binding, and the PQQ β-propeller (Fig. 2A). They may also be further fused to other catalytic domains, such as the ZU5 autopeptidase, which is widely represented in proteolytically processed cell-surface proteins, and the phosphodiester glycosidase (Pfam: NAGPA).

**Figure 2.**
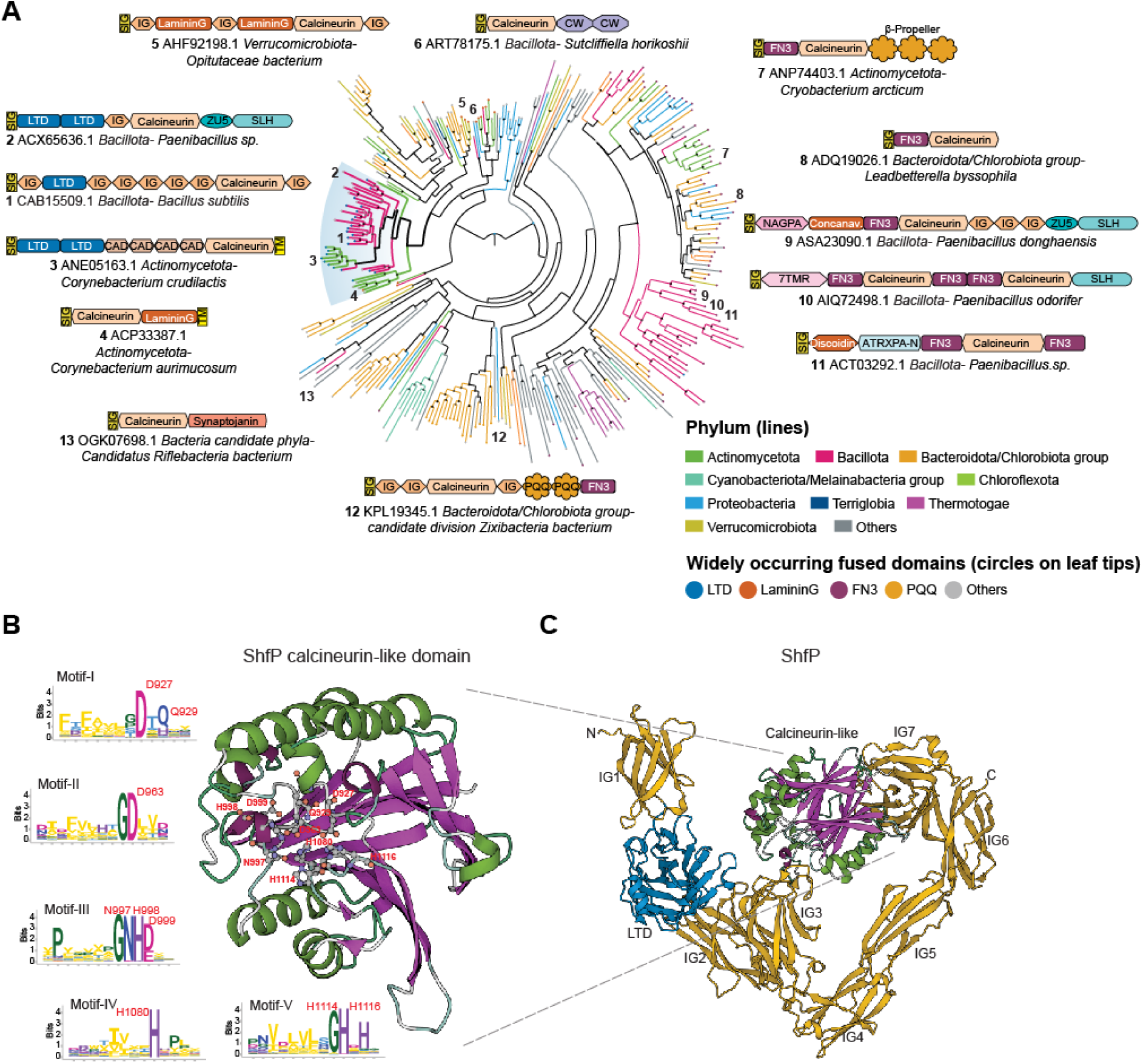
Phylogenetic and domain analysis of ShfP. (A) A phylogenetic tree of a representative set of extracellular calcineurin-like domains from the homologs of the ShfP family is shown. Select domain architectures of proteins are shown near the branch of the tree in which they occur. Numbers indicate the location on the phylogenetic tree of the selected architectures. The branches are colored according to the phylum as shown in the legend; the branch with the *B. subtilis* ShfP and its orthologs is highlighted and shown with thicker lines. The color of the circles at the end of each branch denotes the most common domain in that branch as indicated in the legend, while a gray circle denotes other domain associations. Bootstrap values are shown as a star at each node as grayscale (black = 100%). PQQ: version of the Beta-Propeller; FN3 shown here is a specialized version mapping on to the Pfam model Pur_ac_phosph_N; CW: Pfam model CW_binding (cell wall binding domain); CAD: cadherin; SIG: signal peptide; TM: transmembrane helix; LTD: lamin tail domain; IG: immunoglobulin domain; 7TMR: 7TM receptors with diverse intracellular signaling modules extracellular domain 1; ZU5: ZU5 peptidase; NAGPA: N-acetylglucosamine-1-phosphodiester alpha-N-acetylglucosaminidase; SLH:S-layer homology domain; ATRXPA-N: Anthrax protective antigen glycosidase N-terminal domain. (B) Alphafold2 predicted structure of the *B. subtilis* ShfP calcineurin-like domain with the active sites shown as ball and stick models and the sequence logos of the five conserved motifs of the calcineurin-like domain derived from a multiple sequence alignment of a representative set of ShfP homologs. Letters represent amino acid abbreviations; the height of each letter represents the bit-score of conservation among homologs of the family. (C) Alphafold2 predicted structure of *B. subtilis* ShfP. Gold: Ig domains; blue: LTD domain; green and purple: calcineurin like domain.

The calcineurin-like superfamily comprises universally distributed phosphoesterase domains that feature a four-layered α/β sandwich fold formed from two repeat units of the IF3-C domain (*22*) (Fig. 2B-C). It displays 5 conserved motifs that bind two divalent metal ions (usually Zn(II) or Fe(II)) that catalyze phosphoester hydrolysis. Members of this superfamily act on a variety of organophosphate substrates, including nucleic acids, nucleotides, phosphate-containing lipid head groups, and phosphorylated serines and threonines in proteins (*22*). Indeed, a *B. subtilis* member of one clade of calcineurin-like phosphoesterases (PhoD) has been previously shown to degrade the phosphodiester linkages in cell surface teichoic acids that feature glycerol phosphate or ribitol phosphate (*23*). Based on this precedent and the multiplicity of fusions to potential carbohydrate, adhesion, and DNA-binding domains, we predict that ShfP is likely to adhere to the cell surface macromolecular matrix and catalyze the hydrolysis of phosphoester linkages in one of its components such as (lipo)teichoic acid, eDNA, or a phospholipid head group.

### ShfP is a forespore intermembrane protein requiring the calcineurin-like and adhesion domains for function

The upstream region of the *shfP* gene harbors a putative binding site for σ^G^, a sporulation-specific transcription factor that is active exclusively in the forespore (Fig. 1D) (*24*). To test if *shfP* is expressed in the forespore during sporulation, the *shfP* upstream sequence was fused to super folder-GFP (*sfGFP*) and sfGFP production was visualized by epifluorescence microscopy. We detected fluorescence signal inside the forespore 4 h following sporulation induction, consistent with the timing and location of σ^G^-mediated activation (Fig. 1E-E’). In addition, deletion of σ^G^ or sporulation sigma factors (σ^F^ and σ^E^) that are activated earlier than σ^G^ abrogated the forespore-specific activation of GFP (Fig. 1F-H’), while deletion of the downstream sigma factor (σ^K^) did not (Fig 1I-I’). Thus, unlike *shfA*, which is expressed earlier in the mother cell by σ^E^ (*17, 18*), *shfP* is a sporulation gene that is expressed later in the forespore by σ^G^. We then fused ShfP to sfGFP and determined its localization during sporulation. Consistent with its predicted signal peptide, 4 h after induction of sporulation, ShfP-sfGFP localized to the forespore periphery in an otherwise WT cell (Fig. 1J-J’), and deletion of *shfA* did not disrupt the localization of ShfP-sfGFP (Fig. 1K-K’).

We next tested if the predicted phosphoesterase activity of ShfP is required for function by disrupting conserved active site residues in the calcineurin-like phosphoesterase domain (Fig. 2B). Substituting D927, D963, NHD997-999, H1080, or H1114 and H1116 with Ala did not diminish steady state levels of ShfP (Fig. S1B) but resulted in a partial or total failure of these variants to complement the Δ*shfP* strain (Fig. 1B, lanes 6-10), suggesting that the predicted ShfP phosphoesterase activity is needed for sporulation inhibition. Additionally, deleting the LTD displayed a partial defect (Fig. 1B, lane 12), consistent with the predicted role of this region in anchoring ShfP to the cell surface.

The localization pattern of ShfP-sfGFP and the presence of a signal peptide (Fig. 2C) led us to wonder if ShfP could reside in the intermembrane macromolecular matrix surrounding the forespore (Fig. 1A, “I.M.”). To discern if ShfP resides between the two membranes, which are spaced ∼50 nm apart, we employed a super-resolution fluorescence microscopy technique called DNA-PAINT (*25*). To visualize membranes, we produced a small membrane-bound amphipathic α-helix fused to mCherry prior to induction of sporulation and detected this fusion using antibodies directed against mCherry (*26*). In strains producing this fusion, we co-produced ShfP fused to GFP and detected the fusion using antibodies directed against GFP. Using this technique, the forespore envelope appeared to have > 2-fold increase in thickness relative to the membrane surrounding the mother cell, consistent with the presence of two membranes (Fig. 1L-M, red), and the ShfP-GFP signal (Fig. 1L-M, cyan) localized as puncta at the center of the fluorescence from the membrane stain at the forespore periphery, consistent with an intermembrane space residence of ShfP. In contrast, SpoIVA-GFP, which is produced in the mother cell and resides on the surface of the outer forespore membrane (*27*), visualized using the same technique, displayed a biased localization away from the center of the membrane fluorescence (Fig. 1N-O). Consistent with the intermembrane localization pattern, production of ShfP that did not harbor a signal peptide failed to complement the deletion of *shfP* (Fig. 1B, lane 11; Fig. S1B). Taken together, we conclude that ShfP is a sporulation-specific phosphoesterase that is produced in the forespore and translocated into the forespore intermembrane macromolecular matrix.

### ShfP produces a diffusible extracellular molecule that inhibits sporulation against which ShfA provides immunity

Spo0A is the master regulator for the entry into sporulation (*28*). To study the effect of the ShfA-ShfP pathway, we monitored progression through sporulation using a strain that produces GFP under control of a promoter (*P*_spoIIE_) that is activated when levels of phosphorylated Spo0A reach an adequate threshold that is needed to drive the sporulation program (*10, 29*). We then quantified GFP fluorescence using flow cytometry to monitor individual cells that had initiated sporulation. In an unsynchronized culture, 63% ± 15% of WT cells displayed Spo0A activation (Fig. 3A, “asynchronous”) in early stationary phase, but Δ*shfA* cells (18% ± 16%) displayed reduced Spo0A activation at this time point, even though *shfA* is only expressed after, and is ultimately dependent on, Spo0A itself. This reduction in Spo0A activation was ameliorated in both Δ*shfP* and Δ*shfA* Δ*shfP* strains (Fig. 3A, “asynchronous”).

**Figure 3.**
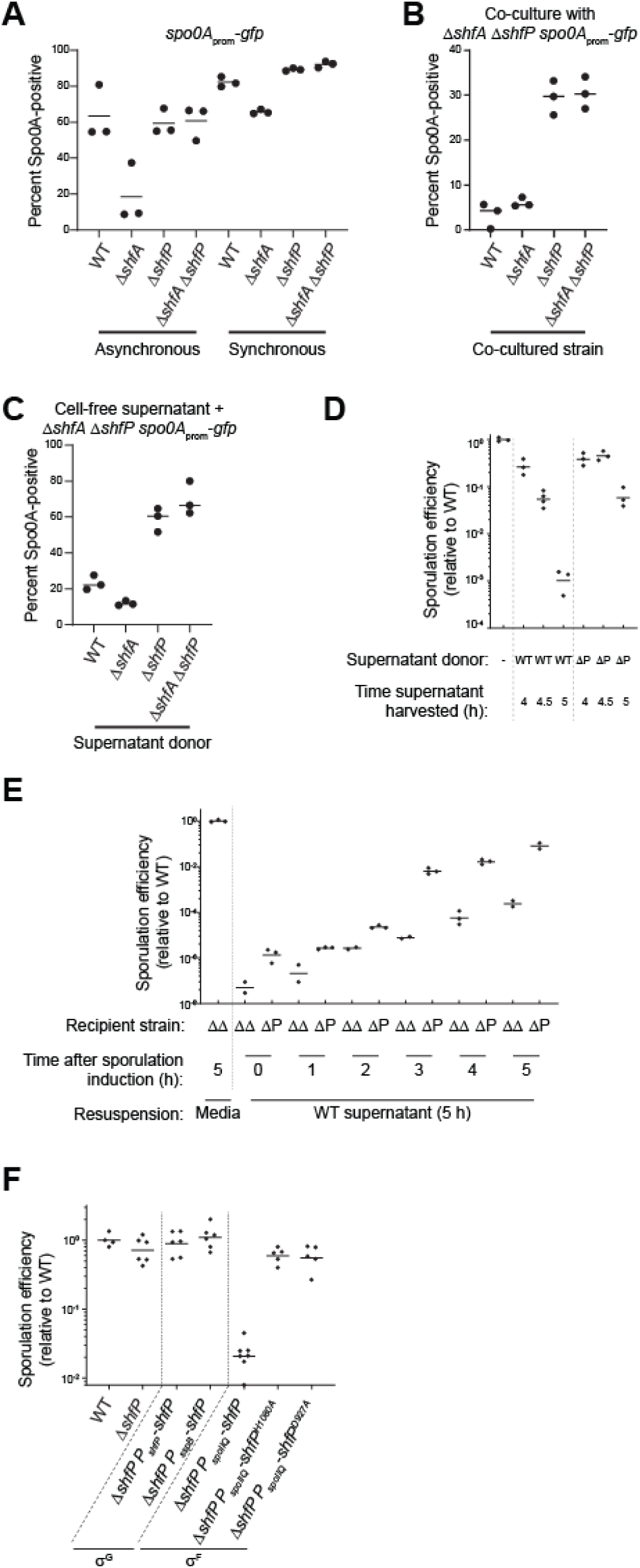
ShfP^+^ cells release an extracellular factor to inhibit late-sporulating cells from entering the sporulation pathway. (A) Percent of Spo0A-positive cells (in cells harboring *P*_spoIIE_-*gfp*: GFP produced under *spo0A* control), determined by flow cytometry, in otherwise WT, Δ*shfA*, Δ*shfP*, or Δ*shfA* Δ*shfP* cells in (left) asynchronous or (right) synchronous cultures. (B-C) Percent of Spo0A-positive Δ*shfA* Δ*shfP* cells harboring the *P*_spoIIE_-*gfp* reporter, determined by flow cytometry, (B) when co-cultured with the indicated strain, or (C) when induced to sporulate with cell-free culture supernatant harvested at t = 5 h after induction of sporulation from the indicated strain. Strains: MF277, CC218, CC219, CC220. (D-E) Sporulation efficiencies of: (D) Δ*shfA* Δ*shfP* cells induced to sporulate in fresh media (lane 1), or cell-free culture supernatant harvested at the indicated times after induction of sporulation from WT (lanes 2-4) or Δ*shfP* cells (lanes 5-7); (E) indicated recipient strains (Δ*shfA* Δ*shfP* or Δ*shfP*) were induced to sporulate for the indicated times, whereupon the cultures were centrifuged and the supernatant replaced with cell-free culture supernatant harvested at the indicated times after induction of sporulation from WT cells. (F) Sporulation efficiency of cells expressing *shfP* under control of σ^G^ (lanes 3-4) or *shfP* and catalytically inactive *shfP* alleles under control of σ^F^ (lane 5-7). Strains: PY79, TD517, TD507, TU36, TU40, TU41, CC134, CC160.

To explain how deletion of a late sporulation gene can affect activation of an early gene in the pathway, we hypothesized that ShfP, produced by cells that entered sporulation early, somehow signals to cells that have not yet entered sporulation to delay their entry into sporulation. In this model, the presence of ShfA in early sporulating (ShfP-producing) cells provides immunity from this ShfP-mediated sporulation delay. Since this model demands asynchronous entry into sporulation, we tested it by repeating the experiment with the Spo0A reporter strains, but instead induced sporulation using the resuspension method (*30*), which synchronizes cells to enter sporulation more uniformly. In a synchronized population of cells, the Spo0A activation defect of the Δ*shfA* strain was largely eliminated (Fig. 3A, “synchronous”), which was consistent with a model in which early sporulating cells actively delay sporulation entry in cells that had not yet activated Spo0A.

To directly test this model, we devised a co-culture system that contained two different strains: Δ*shfA* Δ*shfP* cells harboring the *P*_spoIIE_-*gfp* reporter to assess the entry into sporulation in a sporulation-competent strain (Fig. 3A, lane 4) that did not harbor the ShfA-ShfP pathway, and different co-cultured strains that could influence the strain harboring the reporter. We then analyzed Spo0A activation using flow cytometry. Two hours past the onset of stationary phase, co-culturing the Δ*shfA* Δ*shfP* cells with strains that produced ShfP (WT, Δ*shfA*) resulted in a Spo0A activation defect (Fig. 3B, lanes 1-2), whereas co-culturing the Δ*shfA* Δ*shfP* cells with strains that did not produce ShfP (Δ*shfP*, Δ*shfA* Δ*shfP*) permitted Spo0A activation in the Δ*shfA* Δ*shfP* reporter strain (Fig. 3B, lanes 3-4). To test if this ShfP influence *in trans* is mediated by a soluble extracellular factor or cell-cell contact, we repeated the experiment using cell-free supernatant produced by different donor strains instead of co-culturing. We harvested cell-free supernatants of cultures of various donor strains 5 h after induction of synchronous sporulation and induced sporulation of the recipient Δ*shfA* Δ*shfP* harboring the *P*_spoIIE_-*gfp* reporter using these supernatants. After 2.5 h, we analyzed Spo0A activation in the recipient strain using flow cytometry. Supernatants harvested from cells producing ShfP (WT, Δ*shfA*) repressed the activation of Spo0A (Fig. 3C, lanes 1-2) relative to those harvested from cells that did not produce ShfP (Δ*shfP*, Δ*shfA* Δ*shfP*; Fig. 3C, lanes 3-4), indicating the presence of a ShfP-produced soluble sporulation-delaying factor.

To test the timing of production of the soluble molecule, we harvested culture supernatants from WT and Δ*shfP* cells at 4 h, 4.5 h, and 5 h after induction of synchronized sporulation, lyophilized the sample, concentrated the material ∼4-fold by resuspending in fresh sporulation medium, induced sporulation of the Δ*shfA* Δ*shfP* recipient strain using this supplemented medium, and measured sporulation efficiency. Cells that were resuspended in material harvested 4 h after induction of either WT or Δ*shfP* exhibited a modest decrease in sporulation (Fig. 3D). However, cells resuspended in material harvested at later time points (coincident with the timing of σ^G^-activated expression of *shfP*), exclusively from WT cells exhibited ∼1000-fold reduction in sporulation. Conversely, we induced synchronous sporulation in Δ*shfA* Δ*shfP* and Δ*shfP* strains and, at different time points, replaced the culture supernatants with those derived from WT (Fig. 3E) or Δ*shfA* Δ*shfP* (Fig. S2A) cells at 5 h (which contains the ShfP generated product in the WT supernatant). At initial time points, addition of the WT supernatant did not alter sporulation efficiency but replacing the supernatant any time after t = 3 h (which coincides with the expression of *shfA*) resulted in decreased sporulation efficiency in the Δ*shfA* Δ*shfP*, but not in the Δ*shfP* strain (which produces ShfA; Fig. 3E). However, addition of culture supernatant from Δ*shfA* Δ*shfP* cells did not alter sporulation efficiencies of the recipient cells at any tested time point (Fig. S2A). Thus, ShfA activity manifests 3 h after induction of sporulation, whereas ShfP activity occurs 5 h after sporulation induction. We next tested the effect of expressing *shfP* before *shfA* by cloning the *shfP* gene under control of an earlier forespore-specific transcription factor (σ^F^). Complementing the Δ*shfP* strain with σ^F^*-* expressed *shfP* (which is prior to *shfA* expression) resulted in reduced sporulation efficiency (Fig. 3F, lane 5), whereas expressing the defective *shfP*^H1080A^ or *shfP*^D927A^ alleles earlier under σ^F^ control (Fig. 3F, lanes 6-7) or expressing *shfP* at the proper time and compartment under control of a different σ^G^ promoter (Fig. 3F, lane 4) did not affect sporulation. The data are therefore consistent with a requirement for the earlier production of ShfA to provide a protective benefit against the deleterious effects of the ShfP-generated diffusible product.

### ShfP generates extracellular glycerol, which delays entry into sporulation

To identify the ShfP-produced diffusible factor, we sought to purify the sporulation-delaying activity. Separation of cell-free culture supernatants generated by either WT or Δ*shfP* cultures using size exclusion chromatography revealed several differences in detected peaks (Fig. 4A, top). One fraction (#19) displayed sporulation inhibition activity only in supernatants derived from WT cultures, but not Δ*shfP* cultures (Fig. 4A, bottom). To determine the structure of the released molecule, we lyophilized the active fraction and employed NMR and mass spectrometry. Analysis of ^1^H and ^13^C NMR spectra, along with 2D correlation spectra, showed signals that could only be assigned to glycerol in this subfraction (Fig. 4B-C), and the mass spectrum showed a molecular weight of 93, consistent with glycerol. Lastly, we compared the spectroscopic data with a glycerol standard. The retention time of the active compound and the standard sample (1 mg/mL glycerol) were the same in the extracted ion chromatogram, with both peaks showing a positive mass ion peak for glycerol (Fig. 4D). Quantification of the amount of extracellular glycerol in the sporulation medium 5 h after induction of sporulation using a coupled enzymatic reaction revealed that WT cells produce 1.22 mM ± 0.07 mM glycerol, whereas Δ*shfP* only produce 0.31 mM ± 0.10 mM glycerol (Fig. 4E). Consistent with the notion that the phosphoesterase activity of ShfP is required for glycerol generation, cells that produced variants of ShfP with substitutions that disrupted the phosphoesterase domain produced similar amounts of glycerol as the Δ*shfP* strain (Fig. 4E). Finally, the addition of 1.2 mM, but not 0.2 mM, glycerol to the sporulation media of Δ*shfA* Δ*shfP* cells resulted in a ∼100-fold sporulation defect (Fig. S2B). Complementation of the Δ*shfA* Δ*shfP* strain with *shfA* at a single chromosomal locus corrected the glycerol-mediated sporulation defect, indicating that *shfA* provides protection against the extracellular glycerol (Fig. S2C). We conclude that cells entering sporulation early generate glycerol in a ShfP-dependent manner, which inhibits and delays entry into sporulation for cells that have not already initiated the program.

**Figure 4.**
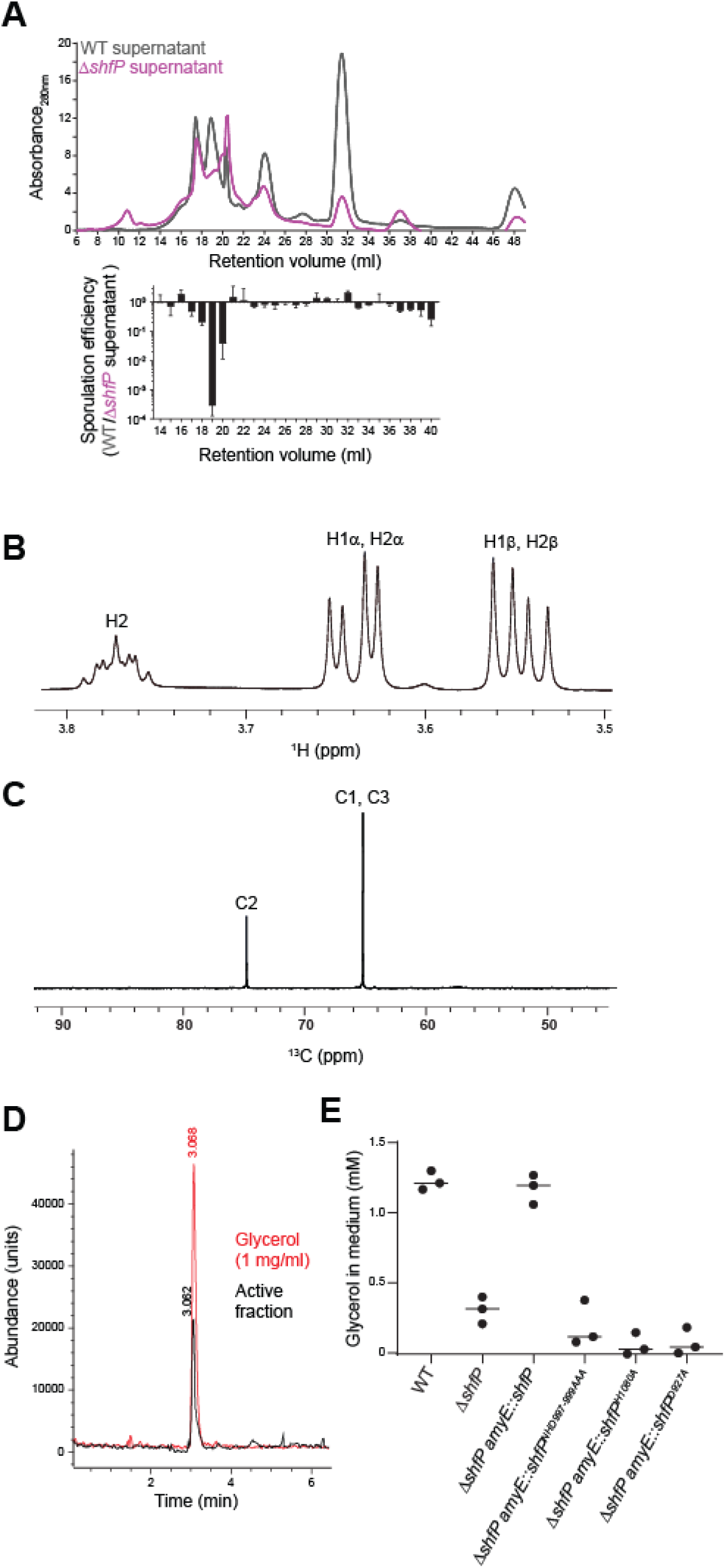
ShfP-dependent release of glycerol by early-sporulating cells inhibits entry into sporulation. (A) Above: size exclusion chromatogram of cell-free culture supernatants harvested from WT (gray) or Δ*shfP* (pink) cells. Below: relative sporulation efficiencies of Δ*shfA* Δ*shfP* cells (strain TD507) subjected to sporulate in cell free culture supernatants harvested from both strains from the above FPLC fractions. Strains: PY79; TD517. (B) ^1^H and (C) ^13^C NMR spectra of active sporulation-inhibitor fraction of purified culture supernatants from WT cells. (D) Extracted ion chromatogram of (black) purified active sporulation-inhibitor fraction and (red) glycerol standard. (E) Concentration of glycerol released into the culture medium of indicated strains. Bars represent mean; data points represent a measure from an independent experiment. Strains: PY79, TD517, CC134, CC254, CC255, CC258.

### The ShfAP pathway promotes heterogeneous entry into sporulation

Although deletion of *shfA* and *shfP* did not result in a sporulation efficiency defect, we hypothesized that inhibiting the entry of a subpopulation of cells into sporulation would affect the heterogeneous entry into the sporulation program. To test this, we observed the fraction of cells initiating sporulation in an asynchronous population using fluorescence microscopy to monitor polar septum formation and forespore engulfment. Over time, WT cells displayed a steady increase in the fraction of cells entering sporulation, but at each time point, we observed a relatively larger fraction of Δ*shfAP* cells that had initiated sporulation (Fig. 5A). Concomitantly, Δ*shfAP* cells also completed sporulation faster relative to WT cells, as evidenced by their elaboration of bright forespores (as viewed by DIC optics, indicating dehydration of the spore core) and mature, released spores (Fig. S3A). To test if the larger proportion of WT cells that had not initiated sporulation confer a growth advantage when faced with an influx of nutrients, we initiated asynchronous sporulation and, at different time points, removed an aliquot of the cells and resuspended them in fresh medium to permit regrowth, and measured initial doubling time. At each time point that we examined, we observed that Δ*shfA*, Δ*shfP*, and Δ*shfAP* cultures displayed a slower doubling time relative to WT, indicating that the absence of the ShfAP pathway resulted in a population that was less poised to take advantage of fresh nutrients after the initiation of sporulation, presumably because these mutants harbored a smaller subpopulation of cells that had not entered sporulation (Fig. 5B, S3B). To directly observe population heterogeneity, we monitored Spo0A activity using flow cytometry at various times after sporulation induction in asynchronous cultures of either WT or Δ*shfAP* cells harboring the *P_spoIIE_-gfp* reporter. Fig. 5C shows the Spo0A activation profiles of three independent cultures of WT (gray) and Δ*shfAP* (red) cells at various time points. Qualitatively, we noticed that the WT cultures displayed multiple populations of cells activating Spo0A, whereas the distribution of Δ*shfAP* cells appeared more Gaussian. This difference in population heterogeneity was reflected in a higher bimodal coefficient (*31*) for WT cells (0.45 ± 0.02) compared to Δ*shfAP* cells (0.37 ± 0.03) at t = 18.5 h. As a further test, we plotted the flow cytometry data for the three independent cultures of WT and Δ*shfAP* cells at t = 18 h as a box plot (Fig. 5D) and examined the distribution of statistical outliers. Cultures of Δ*shfAP* cells showed a roughly equal distribution of statistical outliers at high and low levels of Spo0A activation, whereas outliers in WT cells were heavily skewed towards low activation of Spo0A, suggesting that WT cells maintained a larger population of cells that had likely not committed to enter the sporulation pathway.

**Figure 5.**
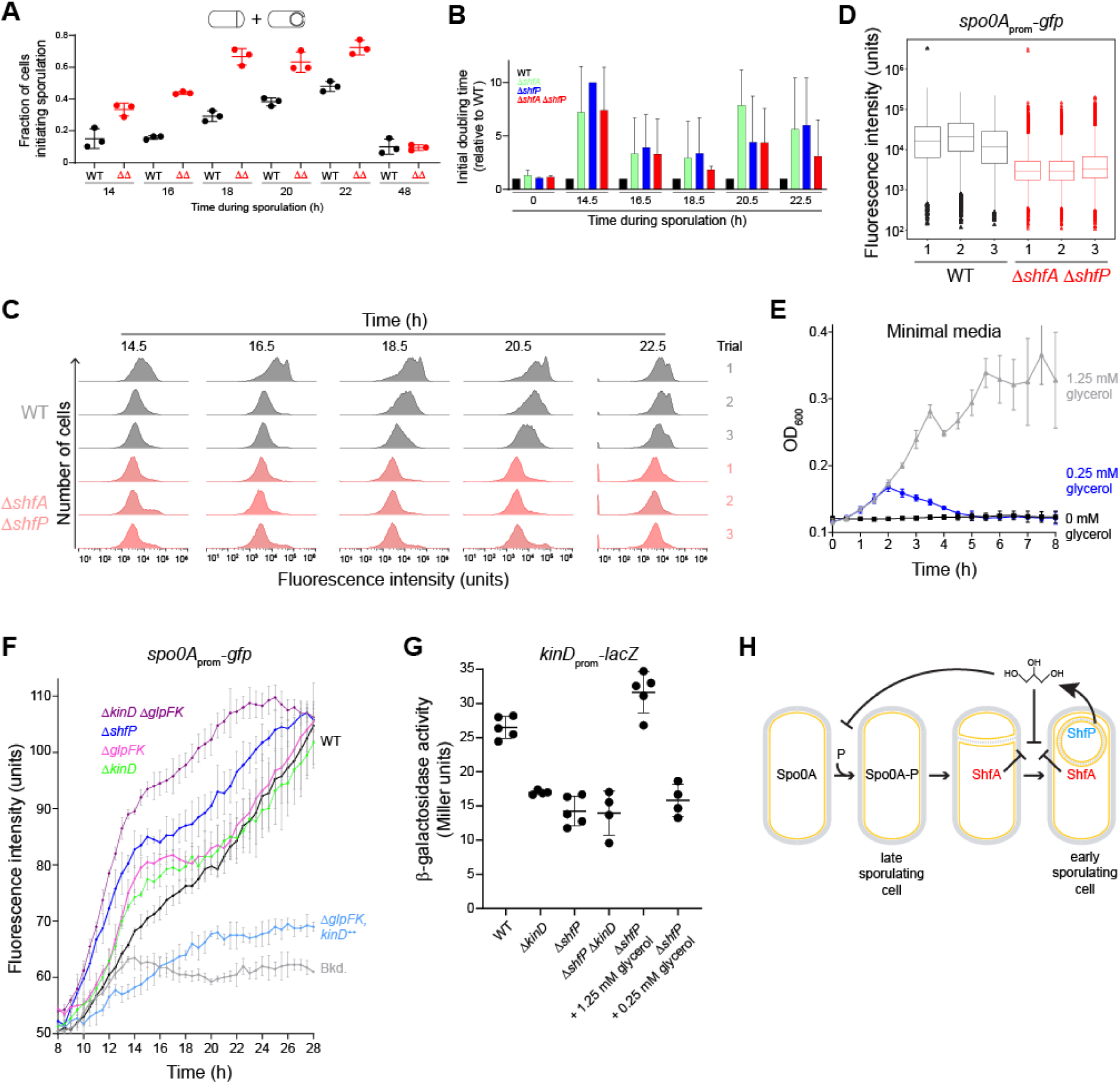
Glycerol secretion enhances phenotypic heterogeneity in sporulating populations. (A) Fraction of (black) WT or (red) Δ*shfA* Δ*shfP* cells committed to sporulation (as evidenced by formation of a polar septum or engulfing membrane using fluorescence microscopy, as shown in the cartoon depiction above) at indicated time points. Bars represent mean; data points represent a measure from an independent experiment. Strains: MF277; CC219. (B) Calculated initial doubling times of indicated strains, relative to WT, induced to sporulate by gradual nutrient deprivation, then resuspended in fresh growth media at indicated time points. Strains: MF277, CC218; CC219; CC220. (C) Histograms of fluorescence intensities in WT and Δ*shfA* Δ*shfP* cells harboring the *P*_spoIIE_-*gfp* reporter induced to sporulate by gradual nutrient deprivation and analyzed at the indicated time points using flow cytometry. Strains: MF277, CC219. (D) 18.5 h time point in (C) represented as box-and-whisker plots. Black: WT; red: Δ*shfA* Δ*shfP*. Box represents interquartile range; bar represents median; whiskers show the range from minimum to lower quartile, and upper quartile to maximum. Statistical outliers are shown as triangle data points. (E) Growth curves, measured by optical density at 600 nm of WT cells grown in minimal media supplemented with (black) 0 mM, (blue) 0.25 mM, or (gray) 1.25 mM glycerol as the sole carbon source. (F) Fluorescence intensity of indicated strains harboring the *P*_spoIIE_-*gfp* reporter induced to sporulate by gradual nutrient deprivation. Strains: PY79; MF277; CC220; TU69; TU78; TU80; TU84. (G) Activation of the KinD-dependent P*_epsA_* promoter in the indicated strains, or in Δ*shfP* cells in the presence of 1.25 mM or 0.25 mM glycerol in the medium. Strains: TU87, TU88, TU89, TU92. (H) Model of early sporulating cells inhibiting Spo0A activation in late sporulating cells. ShfP (blue) generates extracellular glycerol that inhibits Spo0A phosphorylation via KinD activation and glycerol metabolism to sustain vegetative growth. Glycerol also inhibits cortex assembly, which is ameliorated by ShfA.

### Extracellular glycerol acts as a nutrient and a signaling molecule

We next sought to understand the mechanism by which extracellular glycerol delays entry into sporulation. We therefore first tested if the amounts of extracellular glycerol generated by WT, but not Δ*shfP*, cells would be sufficient to sustain growth. In minimal media, addition of 1.25 mM glycerol (similar to the amount generated by WT cells) as the sole carbon source promoted the growth of WT cells (Fig. 5E). In contrast, 0.25 mM glycerol (similar to the amount of glycerol measured in Δ*shfP* cultures) was insufficient to promote growth, indicating that glycerol generated by sporulating cells can serve as a nutrient to delay uncommitted cells from entering the sporulation pathway. Consistent with this model, deletion of two genes involved in glycerol uptake and utilization (*glpF* and *glpK*, respectively) resulted in a slightly faster activation of Spo0A compared to WT (Fig. 5F; compare black and pink traces). However, deletion of *glpF* and *glpK* did not phenocopy the Spo0A activation kinetics of a Δ*shfP* mutant (Fig. 5F, blue trace), suggesting that extracellular glycerol may additionally act to delay sporulation by another pathway.

A previous report indicated that the addition of glycerol to laboratory growth media inhibits Spo0A activation via the signaling histidine kinase KinD with MCP-N and Cache sensor domains (*32*), but the physiological relevance of externally added glycerol was unclear (*33*). We therefore tested if the deletion of *kinD* would result in the faster activation of Spo0A relative to WT. Similar to the deletion of *glpF* and *glpK*, Δ*kinD* cells activated Spo0A faster than WT, but again not at a level comparable to Δ*shfP* cells (Fig. 5F, green trace). However, when we deleted both *glpF* and *glpK*, and additionally *kinD*, Spo0A activation kinetics more closely mimicked that of the Δ*shfP* strain (Fig. 5F, purple trace). In contrast, overproduction of KinD inhibited Spo0A activation (Fig. 5F, blue trace). Finally, we examined the activation of a KinD-dependent gene (*epsA*) in the presence and absence of the ShfAP pathway. Whereas WT cells displayed activation of the *epsA* promoter, deletion of *shfP* failed to activate the promoter, similar to the deletion of *kinD* (Fig. 5G, lanes 1-3). However, external addition of 1.25 mM glycerol (similar to the amount of released glycerol produced by ShfP) activated the KinD-dependent gene, whereas 0.25 mM glycerol (similar to the amount of glycerol present in media of the Δ*shfP* strain) did not (Fig. 5G, lanes 5-6). Consistent with the activation of KinD, addition of 1.25 mM glycerol to a culture of Δ*shfA* Δ*shfP* cells grown asynchronously slowed the entry into sporulation compared to addition of 0.25 mM glycerol (Fig. S3C). KinD contains an MCP-N (a coiled coil region) followed by two Cache domains in its extracellular region. Cache domains are dedicated sensor domains, versions of which bind different diffusible ligands such as citrate (*32*), suggesting that the Cache domains of KinD may act as a glycerol receptor to trigger the intracellular signaling module upon binding it. In sum, the results suggest that ShfP phosphoesterase activity produces extracellular glycerol in sporulating *B. subtilis* cells. This glycerol acts as a signaling molecule to activate KinD to delay sporulation in cells that have not yet committed to enter the sporulation pathway and additionally as a nutrient to delay entry into sporulation to increase the heterogeneity of a differentiating population of cells.

## DISCUSSION

Phenotypic heterogeneity in a clonal population is typically attributed to some inherent randomness: either in gene expression levels that vary between cells, differing quantities of factors that are inherited upon cell division, or changes in the microenvironment that leads to differing cellular responses (*1*). In more extreme cases, ephemeral, but specific, chromosomal alterations occur randomly to switch a cell’s behavior between different states: a phenomenon termed “phase variation”, which can result in heterogeneous behavior (*34*). In this report, we describe a genetically encoded mechanism that directs the programmed generation of glycerol by a subset of sporulating bacteria once those cells have reached a developmental milestone. This extracellular glycerol performs two functions. First, glycerol serves as a nutrient for cells that have not yet entered sporulation, which promotes continued vegetative growth and avoidance of sporulation in those target cells. Second, the released glycerol is a signaling molecule that acts through a sensor kinase (KinD) to actively delay cells from entering the sporulation program. Together, both activities of the released glycerol actively enhance the heterogeneity of the unicellular population and promote the reservation of a subset of cells that do not enter the sporulation program. We observed that, as a result, populations harboring this pathway were better poised to take advantage of a sudden influx of nutrients compared to populations that generated heterogeneity using stochastic gene expression alone. Thus, the genetically encoded nature of this system provided a rare opportunity to directly test the growth advantage of a proposed bet-hedging strategy in a population of clonal unicellular organisms. This altruistic mechanism stands in contrast to the antagonistic cannibalistic behavior that *B. subtilis* exhibits before the onset of sporulation, wherein a subpopulation of cells actively kill their kin to release nutrients to delay their own entry into sporulation (*35*).

*B. subtilis* encodes ∼600 genes, which comprise ∼15% of the genome, that are exclusively upregulated during sporulation. Despite decades of research on sporulation, approximately half of these genes, which included *shfA* (*yabQ*) and *shfP* (*yvnB*), encode for proteins of entirely unknown function. The wide conservation of the *shfA* gene among sporulating Firmicutes suggests that immunity against an extracellular sporulation-delaying molecule may be a shared mechanism for establishing a bet-hedging strategy that is a hallmark of this developmental program in multiple species. In contrast, the ShfP-like phosphoesterases, generating the released glycerol, present a more complex phyletic and potentially functional picture. ShfP-like cell surface phosphoesterases are widely distributed in bacteria and show no particular correlation with the presence or absence of sporulation. Thus, this activity appears to have been coopted for a sporulation-specific function (the generation of glycerol) from what might be a more widespread phosphoesterase function acting on different extracellular macromolecules with phosphoester linkages.

Glycerol was previously implicated in promoting biofilm formation in *B. subtilis*, but this phenomenon was attributed to the use of laboratory growth medium that required glycerol (*33*). We propose that this extracellular glycerol is self-generated, not an intrinsic feature of the environment that *B. subtilis* occupies. The source of the glycerol is not yet clear, but the subcellular localization of ShfP suggests that the substrate for ShfP is likely lipoteichoic acid (whose backbone is phosphoglycerol and therefore could be subject to enzymatic cleavage by the phosphoesterase activity of ShfP) that may be abundant in the intramembrane matrix of the developing forespore. Indeed, early studies have shown that, as *B. subtills* progresses through the sporulation program, lipoteichoic acid is degraded to its monomeric form (*36*). Interestingly, this form was devoid of phosphate, suggesting a concerted action of a phosphodiesterase and alkaline phosphatase activity.

Other studies have pointed to teichoic acids as phosphate stores in *B. subtills* that can be degraded by cell-surface phosphoesterases during phosphate-starvation (*23*). The two phosphoesterases characterized in this process are GlpQ and PhoD and disruption of their genes resulted in a faster vegetative to sporulation transition, evidently because they were unable to prolong post-phosphate-depletion growth. Of these, the teichoate exo-hydrolase GlpQ is a phospholipase C-like TIM barrel enzyme, whereas PhoD, the endo-hydrolase, like ShfP belongs to the calcineurin-like superfamily. Further, analogous to members of the ShfP-related calcineurin-like phosphoesterase radiation, the catalytic domain of PhoD is fused to a FN3 adhesion domain at the N-terminus. This suggests that both PhoD and ShfP might adhere to the cell surface using their respective adhesion domains to process (lipo)teichoate and its derivatives, respectively, during vegetative growth and sporulation.

In light of this, we suggest that the liberation of phosphate from teichoic acid may be used to drive the approximately two-fold increase in ATP production that occurs during sporulation, or even the 30-fold increase in the alarmone nucleotides ppGpp and pGpp (*37*), since it is estimated that up to 40% of total cellular phosphorous is present in lipoteichoic acid (*38*). Another possibility for the source of the glycerol is the head group of the lipid phosphatidylglycerol, which is connected to its acyl chain via a phosphoester bond and would be accessible to the intermembrane space. However, this may be unlikely since, instead of being liberated from phospholipids, glycerol-3-phosphate is instead actively incorporated into phospholipids early in sporulation (*39*). While extracellular glycerol serves to delay sporulation in cells that have not yet entered the pathway, it likely also harbors a toxicity towards cells that produce it. Hence, the glycerol-releasing cells appear to require an “antidote” protein, ShfA, which protects these cells from the evidently toxic effects of glycerol on completing sporulation (Fig. 5H).

The observation that a differentiating population of cells can actively modulate its own heterogeneous behavior through cell-cell communication suggests that the active generation of heterogeneity may be a common phenomenon during development. The communication aspect of this mechanism also raises the possibility that other species or cell types in the immediate environment may be influenced by the released molecule or may even participate in influencing this communication (*40*). Finally, given the likelihood that cell-cell communicated inhibition of sporulation may be a common feature of endospore forming bacteria, it is tempting to speculate that this pathway may be exploited by treating sporulating cells with the diffusible molecule for use as a spore remediation strategy.

## ACKNOWLEDGEMENTS

We thank members of the K.S.R. lab for suggestions and comments on the manuscript; S. Gottesman, G. Storz, A. Khare, V.T. Lee, and S. Wickner for discussions; C. Weiss, E. Kim, M. Tisza, and J. Barriga for strains; F. Soheilian and C. Burks (CCR Electron Microscopy Laboratory) for TEM sample preparation and imaging; the CCR Genomics Core Facility for whole genome sequencing; and R.D. O’Connor for NMR support.

## Author Contributions

K.S.R. supervised the project; K.S.R., T.B.U., T.D., C.C., T.N., and L.A. designed the experiments; T.B.U., T.D., V.A., H.C., C.C., T.N., L.J., L.Z., Y.S., and J.C. performed experiments; T.B.U., T.D., V.A., H.C., C.C., T.N., H.C.-W., L.J., Y.S., H.S., J.C., C.A.B., L.A., and K.S.R. analyzed data; T.B.U., L.A., and K.S.R. wrote the paper.

## Data availability

All data are available in the main text or supplementary materials.

## EXPERIMENTAL PROCEDURES

### Strain construction

Strains used in this study (Table S1) are otherwise isogenic derivates of the *B. subtilis* PY79 strain (*41*). Genes of interest were PCR amplified to include either their native or heterologous promoter and cloned using Gibson Assembly kit (NEB) into integration vectors pDG1662 (for insertion into the *amyE* locus), pDG1731 (for insertion into the *thrC* locus) or pSac-Cm (for insertion into the *sacA* locus) (*42, 43*) containing the appropriate inserts as templates. Site-directed mutagenesis to generate *shfP* mutants was achieved using the QuikChange kit (Agilent). All plasmids were integrated into the *B. subtilis* chromosome by double recombination events at the specified ectopic locus. Plasmid construction was verified by DNA sequencing prior to chromosomal integration.

### Growth conditions for sporulation

*B. subtilis* PY79 or isogenic mutant derivatives were grown on LB plates (10 g Tryptone, 10 g Yeast Extract,10 g Sodium Chloride per liter; KD Medical) with appropriate antibiotic for single colony isolation. Culture conditions for promoting sporulation were done as previously described (*44*). Briefly, for asynchronous sporulation assay, isolated colonies were used to inoculate 2 ml Difco Sporulation Media (DSM, KD Medical) and grown at 37 °C, rotating at 250 rpm, for spore formation. Assays for measuring sporulation efficiencies (see below) were conducted after 24 h incubation, while assays for measuring Spo0A activation, electron microscopy, etc. were done at the indicated times. For synchronous sporulation assay (*30*), isolated colonies were used to inoculate 2 ml casein hydrolysate media (CH KD Medical) and grown overnight at 22 °C rotating at 250 rpm. Overnight cultures were back diluted 1:20 into 20 ml CH media (to OD_600_ ∼ 0.1) and grown 2 h at 37 °C rotating at 250 rpm. Cell cultures were harvested by centrifugation (typically ∼0.9 ml) and pellets resuspended in 1 ml resuspension media (A+B KD Medical) supplemented with 80 μg ml^-1^ threonine (Sigma) for strains lacking a functional *thrC* locus, with 1 mM IPTG if strain contains the inducible P*_hyperspank_* promoter, or the indicated concentration of glycerol. Cells were subsequently grown at 37 °C rotating at 250 rpm and harvested after 24 h for sporulation efficiency assay, or at the indicated time for other assays.

### Sporulation efficiency assay

Sporulation efficiencies were assessed as previously described (*45*). Briefly, WT and mutant *B. subtilis* cells were grown synchronously (in resuspension media) or asynchronously (in DMS) for at least 24 h at 37 °C as described above. Cultures (typically 1- or 2-ml volume) were then exposed to 80°C for 30 min to kill non-sporulating cells. Surviving cells were enumerated by 1/10 serial dilution and plating on LB agar. Viable spores were counted as colony forming units (CFUs); sporulation efficiencies were reported as a ratio to CFUs recovered from a parallel experiment using WT *B. subtilis*. Spontaneous suppressor mutants were isolated by enriching for colonies that grew after multiple rounds of heat-treatment and growth; mutations were identified by whole genome sequencing as described previously (*19*).

### Epifluorescence microscopy

Fluorescence microscopic images of WT and mutant *B. subtilis* were taken as previously described (*46*). Briefly, *B. subtilis* cultures were grown under synchronous or asynchronous sporulation conditions and cells were harvested and resuspended in PBS (KD Medical) at the indicated times. Cells were then stained with 1 ug/ml FM4-64 (Invitrogen) to visualize membranes, then placed on lysine-coated glass bottom dish (MatTek Corp.) under a 1% agarose pad. Cells were viewed with a DeltaVision Core microscope system (Applied Precision) equipped with an environmental control chamber. Images were captured with a Photometrics CoolSnap HQ2 camera. Seventeen planes were acquired every 0.2 μm at 22 °C, and the data were deconvolved using SoftWorx software (GE Healthcare). Control experiments with sporulating strains that did not harbor a *gfp* fusion indicated that the level of GFP fluorescence was well above the limited background fluorescence of the cells.

### Computational analysis of ShfA and ShfP

Homologs for *shfA* and *shfP* were identified using sequence profile searches performed with the PSI-BLAST program (*47*). The searches were run against the non-redundant (NR) protein database of National Center for Biotechnology Information (NCBI), or the sample database compressed by clustering at 50% (nr50) or a custom database of 4210 compete prokaryotic proteomes. Profile-profile searches were conducted using HHpred program (*48, 49*) with multiple alignments to derive the query hidden Markov model augmented by hits against the nr50 or nr70 databases. They were run against 1) PDB; 2) Pfam; and 3) an in-house collection of profiles; the significance of the hits was assessed using the probability percentage and p-value of the HHpred hits. Structure inference was performed using Alphafold2 (*50*). Domain architectures were obtained using a combination of profile searches against Pfam and in-house profiles. Detection of homologs for phyletic pattern correlation analysis was done on a curated set of 4210 complete prokaryotic proteomes. Multiple sequence alignment was built using FAMSA (*51*) and MAFFT (*52*) programs and manually corrected based on secondary structure inferred using Alphafold2 (*50*). Phylogenetic analysis was done using the Le-Gascuel 2008 model in FastTree (*53*). Phylogenetic trees were rendered with the TreeViewer (*54*). Structural visualization of the pdb files were carried out using the Mol*viewer (*55*). The membrane topology was established using deep-learning with the DeepTMHMM program (*56*) and visualized using MembraneFold (*57*).

### DNA-PAINT

No. 1.5 coverslips were first coated with poly-L-lysine (PLL; Sigma) at room temperature for 10 min. The coverslips were then incubated with FluoSpheres beads (0.1 µm, 1:10^6^ dilution; Life Technologies) for 10 min as fiducial markers. *B. subtilis* cells were fixed with 4% Paraformaldehyde (PFA; Thermo Scientific), 0.02% Glutaraldehyde directly in the resuspension buffer for 15 min at room temperature, followed by 30 min incubation on ice. The cells were then washed 3 times with PBS and then resuspend in GTE buffer (50 mM glucose, 20 mM Tris-HCl, 10 mM EDTA). Cells were then incubated with 2 mg ml^-1^ lysozyme at 37 °C for 3 min. The cells were then deposited onto PLL-coated coverslips and incubated for 3 min. After incubation, all the liquid from the sample was removed and the coverslip completely dried at room temperature. The coverslip was then treated with cold (−20 °C) methanol for 5 min and followed by −20 °C acetone for 30 s. The sample was blocked with blocking buffer (2% BSA, 0.1% Triton-x-100 in 1x phosphate-buffered saline) for 30 min at room temperature. Following blocking, the sample was then incubated with single domain anti-GFP (1:300 dilution) antibody and anti-RFP antibody (1:300) conjugated with docking site overnight (Massive Photonics, Germany). The sample was then washed three times with washing buffer (Massive Photonics, Germany). 100 pM imager strands corresponding to different docking site were added to the sample sequentially for two-color DNA-PAINT super resolution imaging. Imaging was performed on a custom-built microscope with a Nikon Ti base and a Nikon N-STORM module equipped with a 100× 1.49 NA oil-immersion lens. Images were acquired with a sCMOS CAMERA (Prime 95B, Teledyne Photometrics) with inclined illumination. A 637 nm laser (Coherent, USA) was used as the excitation wavelength. Typically, 7000 frames were collected with exposure time of 250 ms for DNA-PAINT experiments. Images were analyzed and rendered in Picasso (0.6.0) (*58*). Fluorescence beads were used for drift correction as well as sequential image registration. For image quantification, the spatial relationship between SpoVM-mCherry (used to mark membranes), ShfP-sfGFP, and SpoIVA-GFP in DNA-PAINTING super resolution image was determined using the same line with line width at 5 pixels drawn in different channels (red and cyan) to obtain the fluorescence intensity as a function of position. The center position for each protein was quantified through 2D Gaussian fitting. The standard deviation (S.D.), controlling the spread or width of the curve, obtained from 2D Gaussian fitting was used define the width of protein distributions. The mean position ShfP, SpoIVA and SpoVM were then normalized through subtracting the corresponding mean position of SpoVM. The graph was then plotted as mean ± S.D. (10 different line profiles from 5 different cells) to show the spatial relationship between SpoVM and ShfP, and SpoVM and SpoIVA. The X-axis in the graph shows the relative position (nm) of ShfP or SpoIVA to SpoVM.

### Immunoblotting

Steady state levels of ShfP variants were assessed via immunoblotting as previously described (*59*). Briefly, *B. subtilis* and isogenic mutant cells were induced to sporulate in resuspension media as described above and after 6 h incubation at 37 °C, cells were harvested and resuspended in 500 μl protoplast buffer (0.5 M sucrose, 10 mM K_2_PO_4_, 20 mM MgCl_2_ and 0.1 mg ml^-1^ lysozyme (Sigma)) and incubated at 37 °C for 30 min with shaking at 300 rpm. Protoplasts were harvested by centrifugation and lysed by resuspension in 200 μl PBS and repeatedly passaged through a 20-gauge hypodermic syringe needle (Covidien). 15 μl of the sample was combined with 5 μl of 4× LDS sample buffer (NuPAGE), separated by SDS-PAGE, and transferred to PVDF membrane (Novex) using iBlot (Invitrogen). Blots were blocked in 5% skim milk (Carnation) in Tris-buffered saline (TBS)/Tween (TBS + 1% Tween 20; Sigma) overnight at 4 °C with gentle shaking. Blots were incubated with rabbit antisera raised against purified ShfP or σ^A^ and detected using anti-rabbit IgG StarBright (Bio-Rad) with a ChemiDoc MP imager (BioRad).

### Transmission electron microscopy

Sporulating cells were visualized by transmission electron microscopy as previously described (*19*) with slight modifications. Briefly, cells were allowed to sporulate synchronously in resuspension media for 5 h. Cells were harvested by centrifugation, washed with PBS, and resuspended in equal volume of 4% formaldehyde, 2% glutaraldehyde, and in 0.1 M cacodylate buffer. Resuspended cells were then added to the top of a step gradient of five different metrizoic concentrations (70%, 60%, 50%, 40% and 30%) and centrifuged at 40,000 x g for 60 min at 4 ^°^C. Spore forming cells were found in the middle layers and collected, washed with water, resuspended in fixative, post fixed using 1% osmium tetroxide solution, then dehydrated sequentially in 35%, 50%, 75%, 95% and 100% ethanol followed by 100% propylene oxide. Cells were infiltrated in an equal volume of 100% propylene oxide and epoxy resin overnight and embedded in pure resin the following day. The epoxy resin was cured at 55 °C for 48 h. The cured block was thin-sectioned and stained in uranyl acetate and lead citrate. The samples were imaged with a Hitachi H7600 TEM equipped with a CCD camera.

### Co-culturing

Both reporter and non-reporter strains were grown in 2 mL of DSM at 37 °C with shaking at 250 rpm for 4 h until OD_600_ ∼0.8. Cultures were combined by back-diluting into 6 mL of fresh DSM with equal cell numbers for a total OD_600_ of 0.01 (OD_600_=0.05 for each strain). The combined cultures were grown overnight at 22 °C shaking at 250 rpm. The cultures were then shifted to 37 °C and grown for 2 h past stationary phase (OD_600_ ∼2.5). Cells were then harvested for flow cytometry analysis to measure Spo0A activation (see below).

### Supernatant harvesting

To collect conditioned cell free supernatant, cultures were grown overnight in CH media at 22 °C shaking at 250 rpm. Cultures were then back diluted 1:20 (to OD_600_ ∼0.1) in CH media and grown at 37 °C for 2 h shaking at 250 rpm. Cultures were centrifuged and resuspended in an equal volume of resuspension media (with added threonine if *thrC* locus disrupted) and incubated at 37 °C for 6 h (or otherwise indicated times). The cultures were then centrifuged for 7 min at 7,000 × *g* and supernatant collected and filtered through a 3,000 Dalton MWCO Amicon Ultra-4 filter (Millipore). Aliquots were stored at −80 °C.

For supernatant swapping assays, a culture of the recipient reporter strain was grown overnight in 20 ml CH media at 22 °C shaking at 250 rpm. The culture was then back diluted 1:20 (final OD_600_ ∼0.1) in CH media and grown for 2 h at 37 °C with 250 rpm shaking until it reached mid-exponential phase (OD_600_=0.4-0.6). The culture was then split into equal volumes and centrifuged for 5 min at 7,000 × *g*. The supernatant was discarded, and the cell pellets were resuspended in conditioned cell-free supernatant. Sporulation was allowed to proceed for 2.25 h after induction, and cells were then harvested for flow cytometry analysis to measure Spo0A activation (see below). To determine sporulation efficiencies, the recipient strains were grown overnight in 2 ml CH media at 22 °C shaking at 250 rpm. The cultures were then back diluted 1:20 (OD_600_ ∼0.1) in CH media and continued to grow at 37 °C for 2 h. Then, 0.8 ml of culture was centrifuged, and the supernatant was removed. Next, 2 ml of frozen supernatant were lyophilized from indicated strains and resuspended in 1 ml fresh resuspension media (supplemented with 80 ug/ml threonine) and used to resuspend harvested recipient strains. Cultures were then incubated at 37 °C for 24 h shaking at 250 rpm and sporulation efficiencies were measured as described above.

### Flow cytometry

To analyze Spo0A activation of the reporter *spo0A-gfp* fusion strains, 160 µl of sporulating cultures were centrifuged and the supernatant was removed. Pellets were resuspended in 1 ml of PBS at pH 7.4 (PBS, KD Medical) to a final concentration of ∼10^6^-10^8^ CFU/ml. The samples were then analyzed using a Sony SA3800 flow cytometer. A minimum of 400,000 cells per sample was analyzed using a 488-nm laser and 530/30 bandpass filter to detect GFP signal.

The following voltage values were used: 200 V for forward scatter, 250 V for side scatter, and 400 V for GFP channel. The median GFP signal was measured for each strain and gates were drawn to encompass the GFP signal using isogenic non *gfp*-fused cells as controls to ascertain the amount of background fluorescence the cells emitted. Data was analyzed using FlowJo flow cytometry software. The percentage of the total population of cells with GFP signal above the background was determined as a proxy for the Spo0A activation. Three replicates of each fusion strain were analyzed using the above method for each condition, and the reported values of these three independent replicates, along with the mean value, is shown. To measure Spo0A repression activity in culture supernatants, three independent overnight cultures of the *spo0A-gfp* and the *ΔshfA ΔshfP spo0A-gfp* reporter strains in LB media were back diluted in 200 μl DSM media to 0.1 OD_600_ in a 96-black walls and clear bottom wells microplate (PerkinElmer). The plate was then incubated at 37 ^°C^ with continuous shaking (600 cpm) using a BioTek Synergy H1 microplate reader to record GFP fluorescence, using the 479 Excitation and 520 Emission settings, and OD_600_ measurements were taken every 0.5 h. At the indicated time points, cultures were paused and 50 µl of each culture was diluted in an equal volume of 4% (w/v) Paraformaldehyde in PBS (ThermoFisher) solution and stored at 4 °C overnight. Flow cytometric analysis was performed on 100 µl of sample from each well in the 96-well plate using the same flow cytometric settings as above.

### FPLC

Cell-free supernatants stored at −80 ^°C^ were thawed on ice and filtered through a 0.22 μm membrane to remove precipitants. Then, 1 ml of supernatant was added to a Superdex 30 (GE Healthcare) size exclusion column equilibrated with buffer (25 mM Tris at pH 8.0, 150 mM NaCl) and separated at 0.25 ml min^-1^ using an AKTA *Pure* instrument (GE Healthcare). One ml fractions were collected, lyophilized, and resuspended in 1 ml resuspension buffer to be used on recipient strains to determine sporulation efficiency.

### Nutrient replenishment and outgrowth assay

Three independent overnight cultures of each strain grown in LB media were back diluted in 200 µl DSM media to 0.1 OD_600_ in a 96 well flat-bottom microplate (PerkinElmer). The plate was then incubated at 37 °C with continuous shaking (600 cpm) using BioTek Synergy H1 microplate reader to record OD_600_ measurements every 0.5 h. At the indicated time points, cultures were paused and either 2 µl of each culture was used to seed 200 µl fresh pre-warmed liquid LB media and cultures were allowed to continue growth for the indicated time; or 100 µl of culture was centrifuged, washed with 100 µl PBS, stained with FM4-64 and imaged using epifluorescence microscopy to determine sporulation stage. The initial doubling time for each subculture was calculated as described previously (*60*) using the equation: (Ln(2)/(Ln(OD600_T2_)- Ln(OD600_T1_/T2-T1)).

Alternatively, three independent overnight cultures of the reporter strain were grown overnight in LB media. The overnight cultures were then back diluted in 200 µl DSM media (supplemented with the indicated amount of glycerol) to OD_600_=0.1 in a 96-well (black wall, clear bottom) microplate (PerkinElmer). The plate was then incubated at 37 °C with continuous shaking (600 cpm) using BioTek Synergy H1 microplate reader to record GFP fluorescence, using the 479 Excitation and 520 Emission settings, and OD_600_ measurements every 0.5 h.

### Glycerol detection

Cell-free supernatants from indicated strains were harvested after 5.5 h of growth at 37 °C in resuspension media as described above. Glycerol content in each supernatant fraction was quantitated using the Abcam Free Glycerol Assay Kit (Fluorometric, High Sensitivity; Ab174092) following the manufacturer’s instructions.

### NMR and Mass spectrometry analysis

Cell-free supernatants were separated by Superdex 30 (GE Healthcare) size exclusion column equilibrated with distilled water and separated at 0.25 ml min^-1^. The active fraction (determined by sporulation repression activity) was lyophilized and diluted with HPLC grade 0.1% TFA in water (200 µL) for MS analysis. LC-MS experiments were performed on an Agilent 1260 HPLC system coupled to a 6130 Single Quadrupole mass detector. Solutions (10 µL) were chromatographed on an XSelect HSS T3 Column (4.6 mm X 250 mm, 100Å, 5 µm) eluting with 100% H_2_O (0.1% TFA) at a flow rate of 1.0 mL/ min over 5 min at 20 °C. For verification, glycerol standards (1, 10 mg/mL, Acros Organics) were used as the standard, and pure H_2_O was used as the blank sample. The NMR spectra were recorded on a Bruker Avance 600 MHz NMR using a TCI 5 mm triple-resonance cryoprobe (Bruker, Germany). NMR samples were prepared in 300 mL of D_2_O (Sigma-Aldrich, St. Louis, MO, USA) using a 5 mm Shigemi tube (BMS-005B; Shigemi Co., Ltd, Tokyo, Japan) to obtain higher sensitivity. The structure of glycerol was elucidated based on the 1D and 2D (HSQC, HMBC, COSY) NMR spectra, and mass spectroscopic data analyses.

## SUPPLEMENTAL INFORMATION

**Figure S1.**
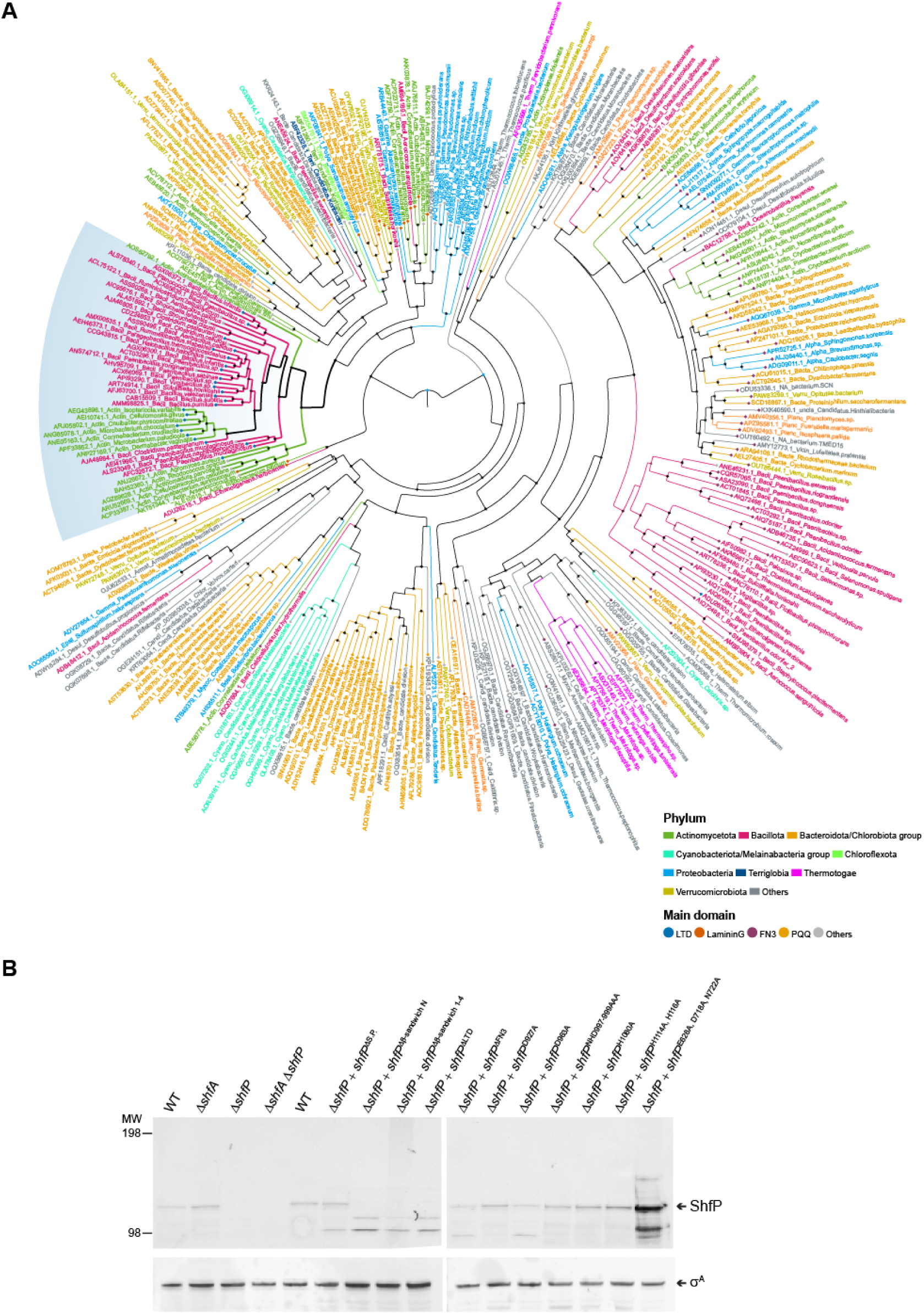
Phylogenetic analysis of ShfP and calcineurin-like proteins. (A) A phylogenetic tree of a representative set of extracellular calcineurin-like domains from the homologs of the ShfP family is shown. The name of each member is shown with its accession number, phylum, and organism name. (B) Western blot against ShfP (top panel) for the following strains: WT (lane 1); Δ*shfA* mutant (lane 2); *ΔshfP* mutant (lane 3); Δ*shfA* Δ*shfP* mutant (lane 4); Δ*shfA ΔshfP amyE::shfP* (lane 5); Δ*shfA ΔshfP amyE::shfP* (*Δ2-28)* (lane 6); Δ*shfA ΔshfP amyE::shfP (Δ27-139)* (lane 7); Δ*shfA ΔshfP amyE::shfP(Δ293-732)* (lane 8); Δ*shfA ΔshfP amyE::shfP(Δ140-280)* (lane 9); Δ*shfA ΔshfP amyE::shfP(Δ1154-1289)* (lane 10); Δ*shfA ΔshfP amyE::shfP(D927A)* (lane 11); Δ*shfA ΔshfP amyE::shfP(D963A)* (lane 12); Δ*shfA ΔshfP amyE::shfP(NHD997-999AAA)* (lane 13); Δ*shfA ΔshfP amyE::shfP(H1080A)* (lane 14); Δ*shfA ΔshfP amyE::shfP(H114A, H116A)* (lane 15); Δ*shfA ΔshfP amyE::shfP(E628A, D718A, N722A)* (lane 16). Anti-sigA used as a loading control (bottom panel). Strains: PY79; CW202; TD517; TD507; CC2; CC19; CC131; CC20; CC138; CC14; CC15; CC16; CC17; CC18; CC166.

**Figure S2.**
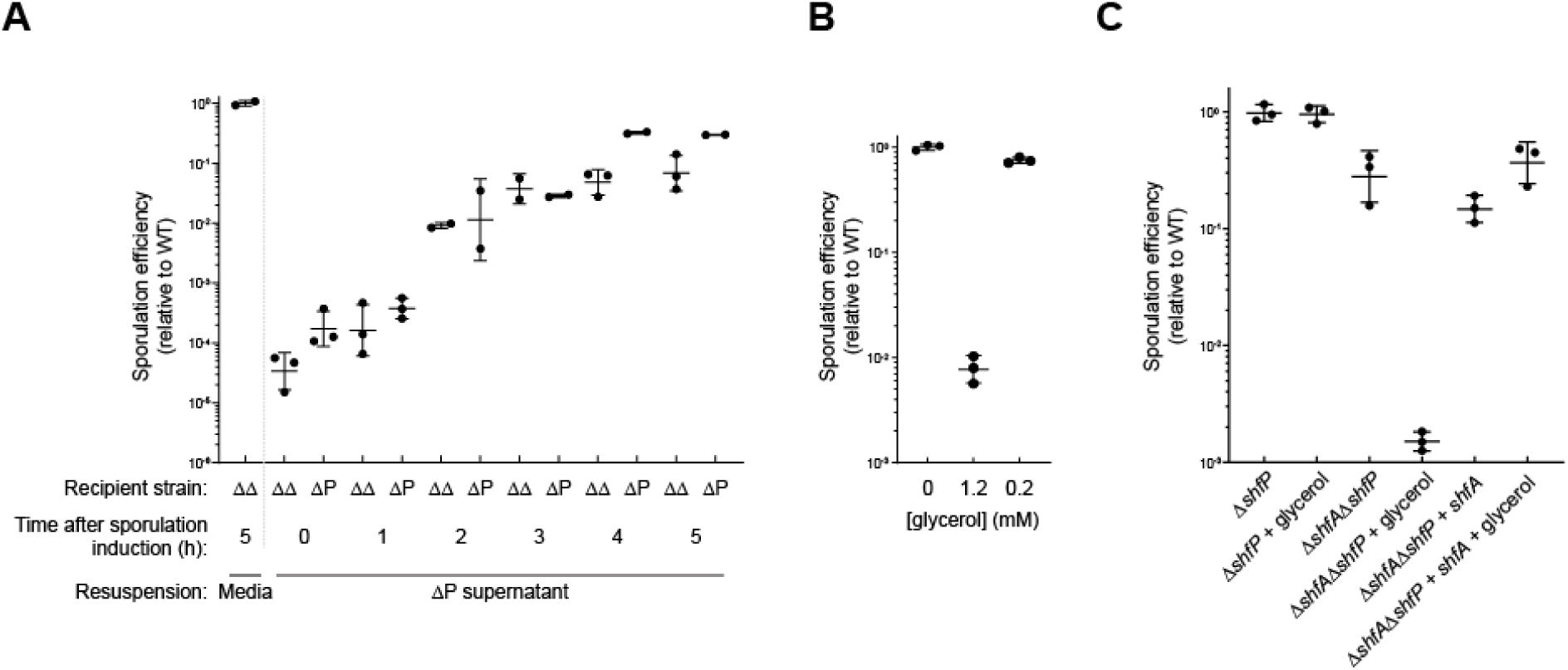
Extracellular glycerol inhibits sporulation. (A) Cell-free supernatant derived from the *ΔshfP* mutant strain show no sporulation suppression activity. Sporulation efficiency of the *ΔshfA ΔshfP* (TD507) and *ΔshfP* (TD517) mutant recipient strains cultured for the indicated times in synchronous sporulation media prior to the addition of cell-free supernatants derived from the *ΔshfP* mutant strain. Sporulation efficiencies are reported as normalized values to that of the recipient strain cultured in the absence of supernatant. Mean and standard deviation of two or more independent cultures for each condition are shown. (B) The addition of glycerol to sporulation media causes a sporulation defect. Average sporulation efficiency of the *ΔshfA ΔshfP* (TD507) mutant strain cultured in synchronous sporulation media with the indicated amount of glycerol added. Sporulation efficiencies are reported as normalized values to that of the recipient strain cultured in the absence of glycerol. Mean and standard deviation of three independent cultures for each condition are shown. (C) Sporulation efficiencies of the *ΔshfP* (TD517), *ΔshfA ΔshfP* (TD507), and *ΔshfA ΔshfP thrC::shfA* (CC221) strains cultured in synchronous sporulation media in the presence or absence of 1.3 mM glycerol. Sporulation efficiencies are reported as normalized values to that of the *ΔshfP* (TD517) recipient strain cultured in the absence of glycerol. Bars represent mean; errors: S.D.

**Figure S3.**
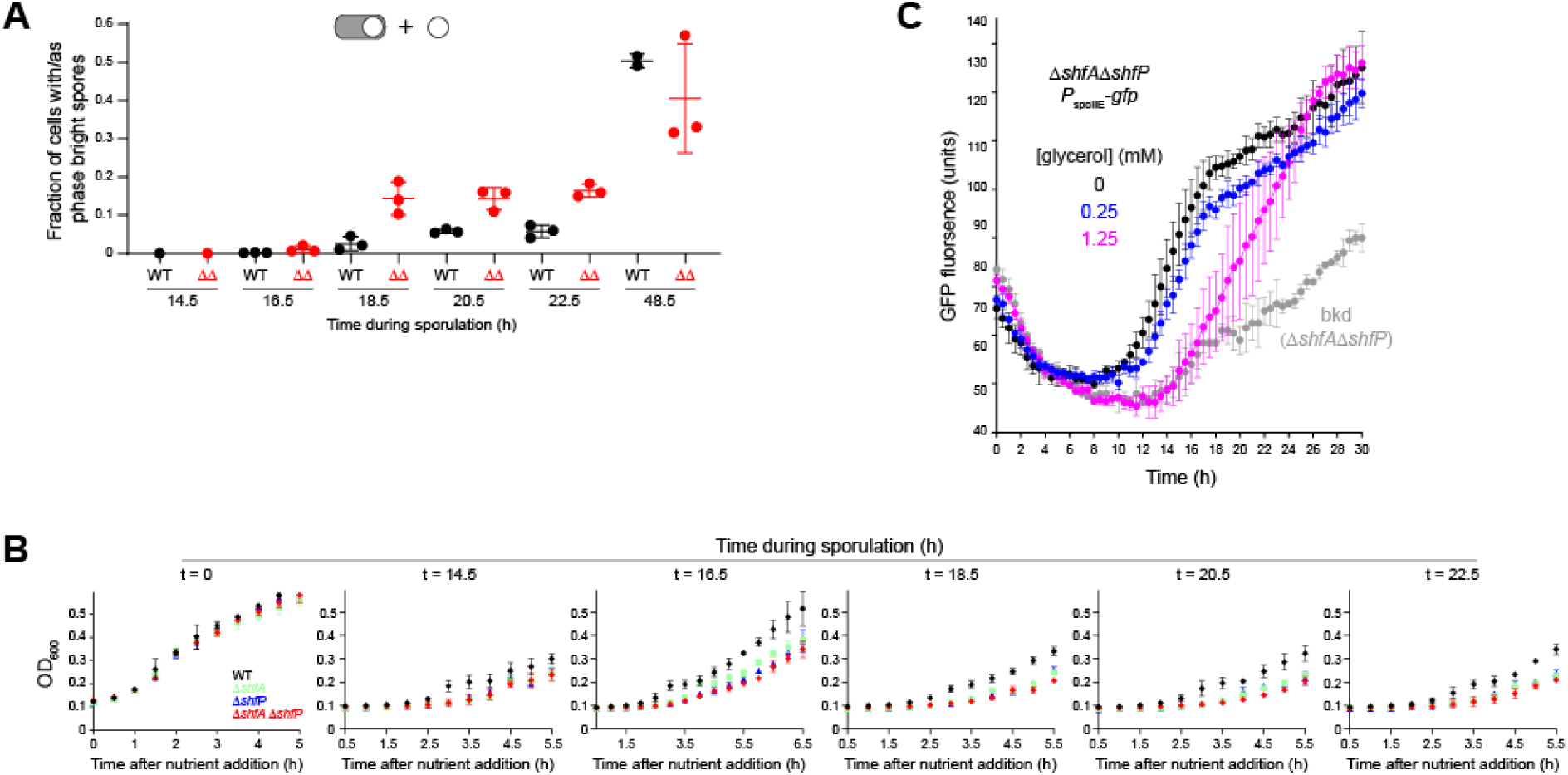
Glycerol secretion delays formation of DIC-bright cells. (A) Fraction of (black) WT (MF277) or (red) Δ*shfA* Δ*shfP* (CC219) cells forming DIC-bright forespore or mature spore structures (as shown in the cartoon depictions above) at the indicated time points. Bars represent mean; errors: S.D.; data points represent a measure from an independent experiment. (B) WT cells respond faster to nutrient replenishment than Δ*shfA* Δ*shfP* mutant cells when grown under asynchronous sporulation conditions. WT (MF277), Δ*shfA* (CC218), Δ*shfP* (CC220) and Δ*shfA* Δ*shfP* (CC219) mutant strains were grown in asynchronous sporulation media. At the indicated time points, 10% of each culture was back diluted in fresh nutrient rich LB media and the OD_600_ was measured over time. Mean and standard deviation of three independent cultures are shown. The initial doubling time for each strain at the indicated time points of subculturing were used for Fig 5B. (C) The addition of glycerol to sporulation media causes a delay in entry into sporulation. The *ΔshfA ΔshfP P_spoIIE_-gfp* (CC219) reporter strain and the non-gfp fusion strain *ΔshfA ΔshfP* (TD507) were grown in synchronous sporulation media with the indicated amount of glycerol added. GFP fluorescence was recorded over time (hours). Mean and standard deviation of four or more independent cultures for each strain and condition are shown.

## REFERENCES

1. A. Sanchez, S. Choubey, J. Kondev, Regulation of noise in gene expression. Annu Rev Biophys 42, 469–491 (2013).

2. I. Levin-Reisman, A. Brauner, I. Ronin, N. Q. Balaban, Epistasis between antibiotic tolerance, persistence, and resistance mutations. Proc Natl Acad Sci U S A 116, 14734–14739 (2019).

3. S. Deshmukh, S. Saini, Phenotypic Heterogeneity in Tumor Progression, and Its Possible Role in the Onset of Cancer. Front Genet 11, 604528 (2020).

4. A. Nguyen, M. Yoshida, H. Goodarzi, S. F. Tavazoie, Highly variable cancer subpopulations that exhibit enhanced transcriptome variability and metastatic fitness. Nat Commun 7, 11246 (2016).

5. J. M. Raser, E. K. O’Shea, Control of stochasticity in eukaryotic gene expression. Science 304, 1811–1814 (2004).

6. L. M. Reyes Ruiz, C. L. Williams, R. Tamayo, Enhancing bacterial survival through phenotypic heterogeneity. PLoS Pathog 16, e1008439 (2020).

7. R. Losick, C. Desplan, Stochasticity and cell fate. Science 320, 65–68 (2008).

8. I. S. Tan, K. S. Ramamurthi, Spore formation in Bacillus subtilis. Environ Microbiol Rep 6, 212–225 (2014).

9. D. Higgins, J. Dworkin, Recent progress in Bacillus subtilis sporulation. FEMS Microbiol Rev 36, 131–148 (2012).

10. M. Fujita, R. Losick, Evidence that entry into sporulation in Bacillus subtilis is governed by a gradual increase in the level and activity of the master regulator Spo0A. Genes Dev 19, 2236–2244 (2005).

11. M. Fujita, J. E. Gonzalez-Pastor, R. Losick, High- and low-threshold genes in the Spo0A regulon of Bacillus subtilis. J Bacteriol 187, 1357–1368 (2005).

12. P. Eswaramoorthy, D. Duan, J. Dinh, A. Dravis, S. N. Devi, M. Fujita, The threshold level of the sensor histidine kinase KinA governs entry into sporulation in Bacillus subtilis. J Bacteriol 192, 3870–3882 (2010).

13. A. Chastanet, D. Vitkup, G. C. Yuan, T. M. Norman, J. S. Liu, R. M. Losick, Broadly heterogeneous activation of the master regulator for sporulation in Bacillus subtilis. Proc Natl Acad Sci U S A 107, 8486–8491 (2010).

14. J. R. Russell, M. T. Cabeen, P. A. Wiggins, J. Paulsson, R. Losick, Noise in a phosphorelay drives stochastic entry into sporulation in Bacillus subtilis. EMBO J 36, 2856–2869 (2017).

15. J. W. Veening, E. J. Stewart, T. W. Berngruber, F. Taddei, O. P. Kuipers, L. W. Hamoen, Bet-hedging and epigenetic inheritance in bacterial cell development. Proc Natl Acad Sci U S A 105, 4393–4398 (2008).

16. C. van Ooij, P. Eichenberger, R. Losick, Dynamic patterns of subcellular protein localization during spore coat morphogenesis in Bacillus subtilis. J Bacteriol 186, 4441–4448 (2004).

17. P. Fawcett, P. Eichenberger, R. Losick, P. Youngman, The transcriptional profile of early to middle sporulation in Bacillus subtilis. Proc Natl Acad Sci U S A 97, 8063–8068 (2000).

18. K. Asai, H. Takamatsu, M. Iwano, T. Kodama, K. Watabe, N. Ogasawara, The *Bacillus subtilis yabQ* gene is essential for formation of the spore cortex. Microbiology (Reading) 147, 919–927 (2001).

19. T. Delerue et al., Bacterial developmental checkpoint that directly monitors cell surface morphogenesis. Dev Cell 57, 344–360 e346 (2022).

20. P. Nicolas et al., Condition-dependent transcriptome reveals high-level regulatory architecture in Bacillus subtilis. Science 335, 1103–1106 (2012).

21. B. J. Mans, V. Anantharaman, L. Aravind, E. V. Koonin, Comparative genomics, evolution and origins of the nuclear envelope and nuclear pore complex. Cell Cycle 3, 1612–1637 (2004).

22. G. G. Nicastro, A. M. Burroughs, L. M. Iyer, L. Aravind, Functionally comparable but evolutionarily distinct nucleotide-targeting effectors help identify conserved paradigms across diverse immune systems. Nucleic Acids Res 51, 11479–11503 (2023).

23. C. L. Myers et al., Identification of Two Phosphate Starvation-induced Wall Teichoic Acid Hydrolases Provides First Insights into the Degradative Pathway of a Key Bacterial Cell Wall Component. J Biol Chem 291, 26066–26082 (2016).

24. S. T. Wang et al., The forespore line of gene expression in Bacillus subtilis. J Mol Biol 358, 16–37 (2006).

25. R. Jungmann, M. S. Avendano, J. B. Woehrstein, M. Dai, W. M. Shih, P. Yin, Multiplexed 3D cellular super-resolution imaging with DNA-PAINT and Exchange-PAINT. Nat Methods 11, 313–318 (2014).

26. E. Y. Kim, E. R. Tyndall, K. C. Huang, F. Tian, K. S. Ramamurthi, Dash-and-Recruit Mechanism Drives Membrane Curvature Recognition by the Small Bacterial Protein SpoVM. Cell Syst 5, 518–526 e513 (2017).

27. E. A. Peluso, T. B. Updegrove, J. Chen, H. Shroff, K. S. Ramamurthi, A 2-dimensional ratchet model describes assembly initiation of a specialized bacterial cell surface. Proc Natl Acad Sci U S A 116, 21789–21799 (2019).

28. D. Schultz, Coordination of cell decisions and promotion of phenotypic diversity in B. subtilis via pulsed behavior of the phosphorelay. Bioessays 38, 440–445 (2016).

29. P. Eswaramoorthy, J. Dinh, D. Duan, O. A. Igoshin, M. Fujita, Single-cell measurement of the levels and distributions of the phosphorelay components in a population of sporulating Bacillus subtilis cells. Microbiology (Reading) 156, 2294–2304 (2010).

30. J. M. Sterlini, J. Mandelstam, Commitment to sporulation in Bacillus subtilis and its relationship to development of actinomycin resistance. Biochem J 113, 29–37 (1969).

31. R. Pfister, K. A. Schwarz, M. Janczyk, R. Dale, J. B. Freeman, Good things peak in pairs: a note on the bimodality coefficient. Front Psychol 4, 700 (2013).

32. V. Anantharaman, L. Aravind, Cache - a signaling domain common to animal Ca(2+)- channel subunits and a class of prokaryotic chemotaxis receptors. Trends Biochem Sci 25, 535–537 (2000).

33. M. Shemesh, Y. Chai, A combination of glycerol and manganese promotes biofilm formation in Bacillus subtilis via histidine kinase KinD signaling. J Bacteriol 195, 2747–2754 (2013).

34. M. Zamora, C. A. Ziegler, P. L. Freddolino, A. J. Wolfe, A Thermosensitive, Phase-Variable Epigenetic Switch: pap Revisited. Microbiol Mol Biol Rev 84, (2020).

35. J. E. Gonzalez-Pastor, E. C. Hobbs, R. Losick, Cannibalism by sporulating bacteria. Science 301, 510–513 (2003).

36. W. Kusser, F. Fiedler, Teichoicase from Bacillus subtilis Marburg. J Bacteriol 155, 302–310 (1983).

37. T. B. Updegrove et al., Reformulation of an extant ATPase active site to mimic ancestral GTPase activity reveals a nucleotide base requirement for function. Elife 10, (2021).

38. Y.-W. Tang, Y.-W. Tang, ScienceDirect, Molecular medical microbiology. (Academic Press, London, ed. Second edition., 2015).

39. M. E. Pedrido, P. de Ona, W. Ramirez, C. Lenini, A. Goni, R. Grau, Spo0A links de novo fatty acid synthesis to sporulation and biofilm development in Bacillus subtilis. Mol Microbiol 87, 348–367 (2013).

40. S. Mukherjee, B. L. Bassler, Bacterial quorum sensing in complex and dynamically changing environments. Nat Rev Microbiol 17, 371–382 (2019).

41. P. Youngman, J. B. Perkins, R. Losick, Construction of a cloning site near one end of Tn917 into which foreign DNA may be inserted without affecting transposition in Bacillus subtilis or expression of the transposon-borne erm gene. Plasmid 12, 1–9 (1984).

42. A. M. Guerout-Fleury, N. Frandsen, P. Stragier, Plasmids for ectopic integration in Bacillus subtilis. Gene 180, 57–61 (1996).

43. R. Middleton, A. Hofmeister, New shuttle vectors for ectopic insertion of genes into Bacillus subtilis. Plasmid 51, 238–245 (2004).

44. S. E. Ebmeier, I. S. Tan, K. R. Clapham, K. S. Ramamurthi, Small proteins link coat and cortex assembly during sporulation in Bacillus subtilis. Mol Microbiol 84, 682–696 (2012).

45. J. P. Castaing, S. Lee, V. Anantharaman, G. E. Ravilious, L. Aravind, K. S. Ramamurthi, An autoinhibitory conformation of the Bacillus subtilis spore coat protein SpoIVA prevents its premature ATP-independent aggregation. FEMS Microbiol Lett 358, 145–153 (2014).

46. P. J. Eswara et al., An essential Staphylococcus aureus cell division protein directly regulates FtsZ dynamics. Elife 7, (2018).

47. S. F. Altschul, E. V. Koonin, Iterated profile searches with PSI-BLAST--a tool for discovery in protein databases. Trends Biochem Sci 23, 444–447 (1998).

48. J. Soding, Protein homology detection by HMM-HMM comparison. Bioinformatics 21, 951–960 (2005).

49. J. Soding, A. Biegert, A. N. Lupas, The HHpred interactive server for protein homology detection and structure prediction. Nucleic Acids Res 33, W244–248 (2005).

50. J. Jumper et al., Highly accurate protein structure prediction with AlphaFold. Nature 596, 583–589 (2021).

51. S. Deorowicz, A. Debudaj-Grabysz, A. Gudys, FAMSA: Fast and accurate multiple sequence alignment of huge protein families. Sci Rep 6, 33964 (2016).

52. K. Katoh, J. Rozewicki, K. D. Yamada, MAFFT online service: multiple sequence alignment, interactive sequence choice and visualization. Brief Bioinform 20, 1160–1166 (2019).

53. M. N. Price, P. S. Dehal, A. P. Arkin, FastTree: computing large minimum evolution trees with profiles instead of a distance matrix. Mol Biol Evol 26, 1641–1650 (2009).

54. G. Bianchini, P. Sanchez-Baracaldo, TreeViewer: Flexible, modular software to visualise and manipulate phylogenetic trees. Ecol Evol 14, e10873 (2024).

55. D. Sehnal et al., Mol* Viewer: modern web app for 3D visualization and analysis of large biomolecular structures. Nucleic Acids Res, (2021).

56. J. Hallgren et al., DeepTMHMM predicts alpha and beta transmembrane proteins using deep neural networks. bioRxiv, (2022).

57. S. Gutierrez, W. G. Tyczynski, W. Boomsma, F. Teufel, O. Winther, MembraneFold: Visualising transmembrane protein structure and topology. bioRxiv, (2022).

58. J. Schnitzbauer, M. T. Strauss, T. Schlichthaerle, F. Schueder, R. Jungmann, Super-resolution microscopy with DNA-PAINT. Nat Protoc 12, 1198–1228 (2017).

59. I. S. Tan, C. A. Weiss, D. L. Popham, K. S. Ramamurthi, A Quality-Control Mechanism Removes Unfit Cells from a Population of Sporulating Bacteria. Dev Cell 34, 682–693 (2015).

60. B. C. Crump, C. S. Hopkinson, M. L. Sogin, J. E. Hobbie, Microbial biogeography along an estuarine salinity gradient: combined influences of bacterial growth and residence time. Appl Environ Microbiol 70, 1494–1505 (2004).

